# Determining the Quantitative Principles of T Cell Response to Antigenic Disparity: The Case of Stem Cell Transplant Donor-Recipient Exome Wide Mismatching and HLA Bound Alloreactive Peptides

**DOI:** 10.1101/305474

**Authors:** Ali Salman, Vishal Koparde, Charles Hall, Max Jameson-Lee, Catherine Roberts, Myrna Serrano, Badar AbdulRazzaq, Masoud Manjili, Dayanjan Wijesinghe, Shahrukh Hashmi, Greg Buck, Rehan Qayyum, Michael Neale, Jason Reed, Amir Toor

## Abstract

Alloreactivity compromising clinical outcomes in stem cell transplantation is observed despite HLA matching of donors and recipients. This has its origin in the variation between the exomes of the two, which provides the basis for minor histocompatibility antigens (mHA). The mHA presented on the HLA class I and II molecules and the ensuing T cell response to these antigens results in graft versus host disease. In this paper, results of a whole exome sequencing study are presented, with resulting alloreactive polymorphic peptides and their HLA class I and HLA class II (DRB1) binding affinity quantified. Large libraries of potentially alloreactive recipient peptides binding both sets of molecules were identified, with HLA-DRB1 presenting an order of magnitude greater number of peptides. These results are used to develop a quantitative framework to understand the immunobiology of transplantation. A tensor-based approach is used to derive the equations needed to determine the alloreactive donor T cell response from the mHA-HLA binding affinity and protein expression data. This approach may be used in future studies to simulate the magnitude of expected donor T cell response and risk for alloreactive complications in HLA matched or mismatched hematopoietic cell and solid organ transplantation.

## Introduction

Graft-versus-host Disease (GVHD) represents a significant cause of morbidity and mortality in stem-cell transplant (SCT) recipients^1^. GVHD in an HLA-matched allogeneic stem cell transplant is the archetype of an adaptive immune response with donor derived T cells responding to recipient antigens presented on shared HLA class I and class II antigens^2, 3, 4^. Since the beginning, HLA matching has been the bedrock principle of donor selection in SCT, and this is particularly so when the donor is not a close relative^5, 6^. Improvements in the fidelity of HLA matching between unrelated transplant donors and recipients has yielded incremental benefits in patient outcomes, with improvements in survival resulting from both a reduction in GVHD risk as well as reduction in graft loss and optimization of relapse risk. Nevertheless, GVHD remains a therapeutic challenge, and there is little that can be done to predict the outcomes of specific donor-recipient pairs.

This challenge may be surmounted by accounting for genomic variation between the donors and recipients which yields the peptides presented on HLA molecules, known as minor histocompatibility antigens (mHA)^7, 8^. While mHA have had a recognized pathophysiologic role in allogeneic SCT outcomes, especially in GVHD pathogenesis, it has not been possible to apply the notion to clinical practice because mHA characterization is a cumbersome process^9, 10, 11, 12^. Two developments in the past decade have changed this situation. One, the emergence of next generation DNA sequencing techniques, such as single nucleotide polymorphism mapping ^13, 14^ and whole exome sequencing (WES) to identify the potential antigenic differences^15, 16^. The second is the development of machine learning algorithms which allow determination of the binding affinity that different antigens may have for specific HLA molecules^17, 18, 19^. These two techniques have been combined to develop algorithms that may be used to determine the complex array of recipient antigens that a given donor T cells may encounter in a recipient^20, 21^. This knowledge of mHA in turn may allow simulation of alloreactive T cell responses in equivalently HLA matched SCT donor-recipient pairs (DRP) to identify donors with optimal alloreactivity.

Studies reporting exome-wide or other genomic disparities in donors and recipients, have demonstrated a large body of DNA sequence differences between transplant donors and recipients, independent of relatedness and HLA matching^14, 15, 16^. These large genomic differences have been translated to peptides and HLA affinities for the resulting peptides determined^20^. This too yields large libraries of antigens which may be analyzed by either simulating alloreactive T cell responses or by statistical methodology to determine predictive power for alloreactive T cell responses^22, 23^. To date, these models have examined recipient peptide presentation on HLA class I and studied the resulting associations.

As noted above, HLA-matched SCT remains fraught with uncertainty as patients with HLA-matched donors continue to have disparate outcomes^24, 25^. A quantitative model of transplant alloreactivity would allow a more complete understanding of the molecular immunology of SCT, help to identify the most suitable donors for specific recipients, and allow personalized determination of the optimal level of immunosuppression. A central assumption in one such quantitative model, the dynamical system model of T cell responses, is that alloreactivity (such as GVHD) risk is a function of the cumulative mHA variation in the context of the HLA type of each donor-recipient pair (DRP), and may thus be regarded as an alloreactivity potential for that pair^15, 20, 26^. Clinical outcomes partially depend on the cumulative donor T cell responses to the burden of polymorphic recipient peptides. Previous work applying this dynamical system model to HLA class I presented molecules demonstrates that there are large differences in the simulated T cell responses between different HLA matched DRP^22, 23^. Herein, previously reported findings of WES of SCT DRP are extended with an analysis of the HLA class II presentation of polymorphic peptides. A comparison of the difference in magnitude of the *derived* peptide libraries presented on the HLA class I and HLA class II molecules in the DRP is presented. Next a mathematical model is developed which may allow the development of a new approach to the study of such large data sets and their eventual application to clinical medicine. The previously reported dynamical systems model of alloreactive T cell responses is generalized to include both HLA class I and HLA class II presented peptides. The model is expanded to account for different conditions T cells may be subject to, specifically their own state of antigen-responsiveness and the cytokine milieu. This quantitative perspective may, in the future, permit successful simulation of alloreactive T cell responses between different donors and recipients in SCT.

## Methods

After obtaining approval from the institutional review board at the Virginia Commonwealth University, whole exome sequencing (WES) was performed on previously cryopreserved DNA samples from 77 HLA-matched DRP (Supplementary Table 1) as previously described^15, 22^. Briefly, whole exome sequences from each DRP were compared with each other, as well as to a standard reference exome. All nonsynonymous single-nucleotide polymorphisms (nsSNPs) present in the recipient and donor were identified and recorded in the vcf format. Subsequent processing of the vcf files was done using custom python scripts to remove synonymous mutations, eliminate duplicates, and record the coordinates of the SNPs. Non-synonymous SNPs that exist in the recipient but not in the donor were recorded and identified as potential source of alloreactive antigens. Non-synonymous, single nucleotide polymorphisms (nsSNP) in each DRP would correspond to potential antigens due to the resulting amino acid substitution in oligopeptides which bind HLA in that DRP (Figure 1A).

**Figure 1.**
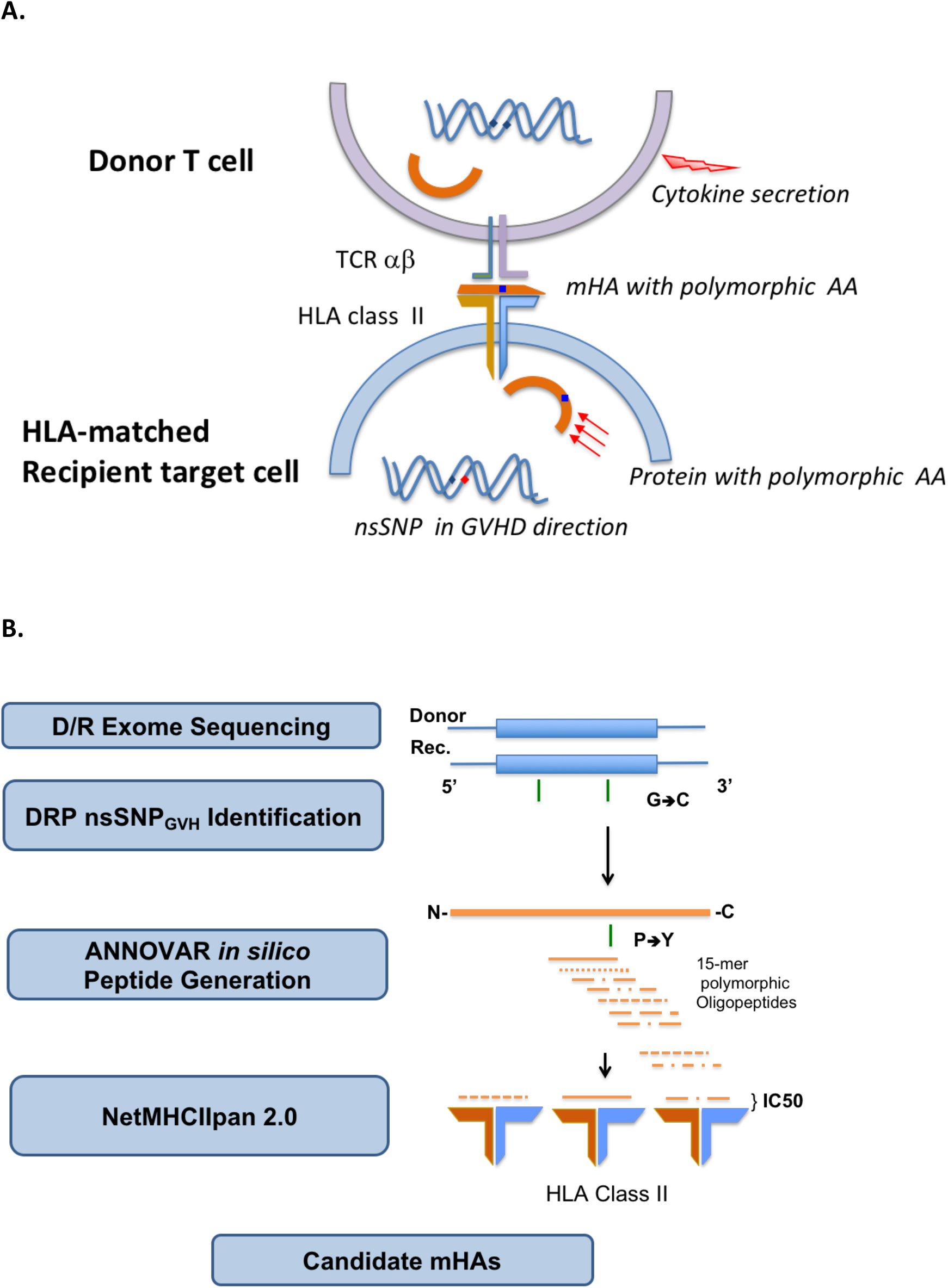
Non-synonymous single nucleotide polymorphisms present in the recipient and absent in the donor yield alloreactive peptides which may be presented to the donor T cells on HLA class I and II molecules. HLA class II presentation and CD4+ T cell recognition and response depicted (A). Schematic depicting the analytic sequence from exome sequencing to HLA class II mHA prediction (B).

To derive the peptide sequences for this study, an average peptide length of 15 amino acids for HLA class II HLA was used^27^. HLA class I bound 9-mer peptides were generated as previously described^20^. Each of the nsSNPs could potentially be incorporated into the alloreactive peptide of 15 amino acids. The position of the nsSNP encoded polymorphic amino acid in the peptide could vary from the N-terminus to the C-terminus of the peptide. The possible library of peptides will thus be contained within a 29-mer oligopeptide (Figure 1B). Thus, there are 15 different HLA-II binding peptides that could potentially be generated from each nsSNP identified by WES. ANNOVAR was used to generate 29-mer peptides for each nsSNP respectively to study HLA class II presentation. In ANNOVAR, a sliding window method was used with the “seq_padding” option of the “annotate_variation” function to generate the 15 different 15-mers resulting from each nsSNP. Tissue expression of the proteins from which the peptides were derived was determined as previously described^23^.

Once the peptide library was created for each DRP, the HLA types for the recipient were tabulated from the medical records. For class II HLA, HLA-DRB1 alleles for each patient were recorded. Each patient’s HLA-DRB1 allele types (and HLA class I alleles, as previously described) along with peptide library were analyzed using NetMHCIIpan 2.0 to derive the binding affinity of each peptide-HLA complex. This was given as an IC50 (half-maximal inhibitory concentration) for each peptide, measured in nano-Molar. This measure of binding affinity provided the concentration of peptide required to displace 50% of a standard peptide from the HLA type to which it would have been bound.

Peptides present in the recipient but absent in the donor, generated from the ANNOVAR sliding window with IC50 values for all the different patient HLA types were tabulated and duplicates were deleted. Any peptide with the same amino acid sequence but different SNP position along the peptide must have generated from a different area of the exome and was therefore retained in the enumeration. When compiling the peptides binding to different HLA alleles, the patients with homozygous allele for DRB1 had their peptide values doubled to simulate having double the normal number of allele-specific HLA bound peptides presented. Analysis of the number of strongly bound (SB; IC50 ≤50 nM) and bound peptides (BP; IC50 ≤500 nM) for each patient-HLA allele combination was done by listing the peptides in descending order of binding affinity, as measured by IC50 levels (Table 1A & 1B). HLA class I and HLA class II bound peptides were compared numerically for this perspective paper.

**Table 1.**
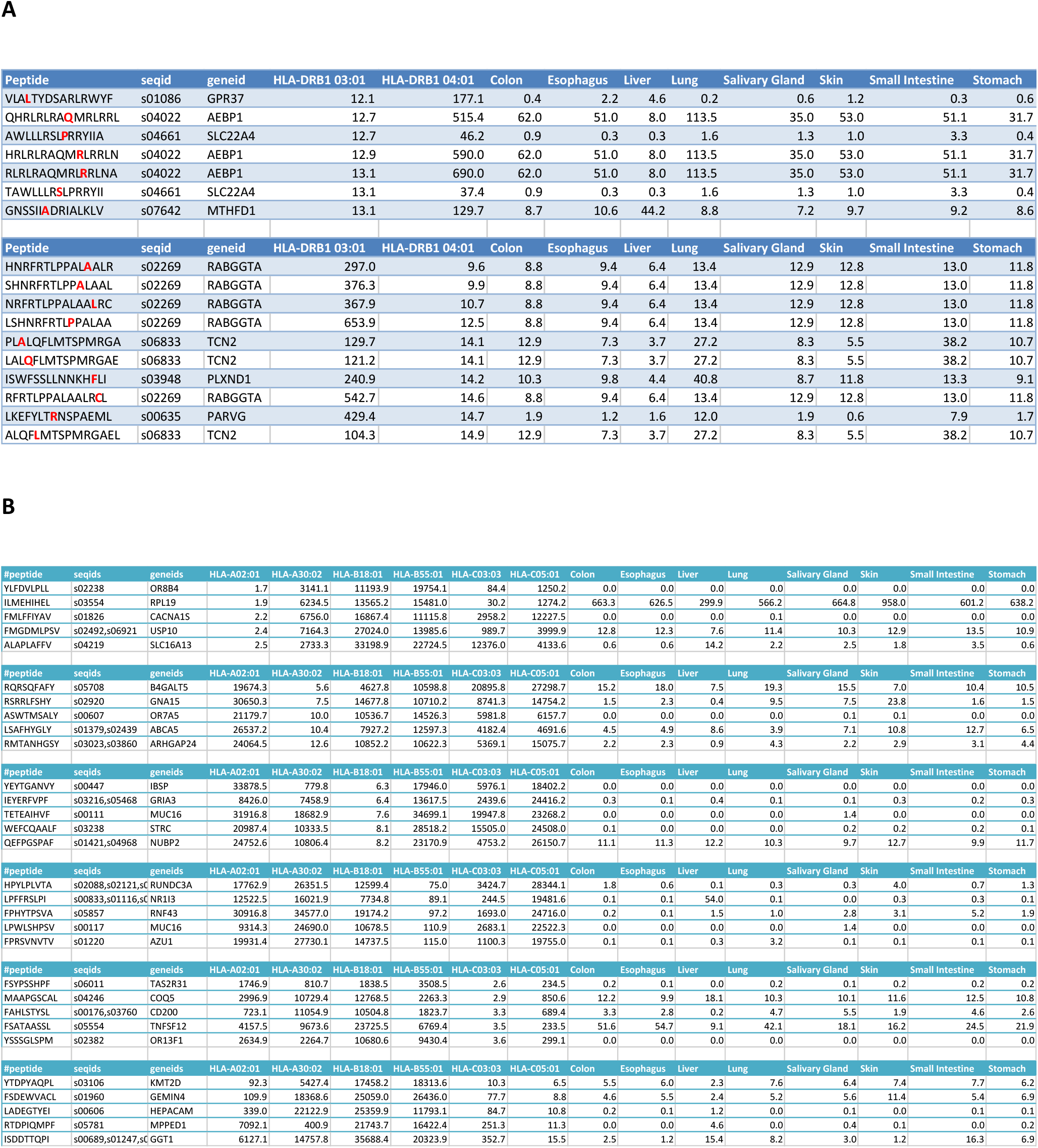
Example of the strong binding peptides for HLA class II (A) and HLA class I (B) in a DRP. Most peptides have relevant interaction with one HLA molecule but some bind multiple HLA molecules, more so with HLA class II molecules.

## Results

### HLA class II bound alloreactive peptides

Whole exome sequencing (WES) was performed on the cohort of 77 donor recipient pairs (DRP) of which 75 were evaluable for this analysis. SNPs were identified, following which alloreactive peptides and HLA-DRB1 binding affinities were derived. HLA matched unrelated donor (MUD) DRP exhibited a higher number of HLA-DRB1-BP; mean: 39,584 alloreactive peptides in HLA matched related donors (MRD) vs. 67,987 in MUD (t-test P <.001). When only the SB peptides are analyzed, this trend while present no longer remains statistically significant, mean SB 6,077 alloreactive peptides in MRD vs. 9,535 in MUD (p=0.168) (Figure 2A & 2B). This is consistent with the larger burden of exome variation in MUD transplant recipients. Significantly more MUD DRP had BP > the median 52,983 peptides for the whole cohort (34/49 vs. 4/26, Fishers Exact test p<0.0001), as well as SB >4,245 (30/49 vs. 8/26, p=0.012), when compared to MRD DRP. There was marked variability in the HLA DRB1 allele binding affinity in the various peptides as well as the tissue expression of the proteins from which peptides were derived (Table 1A). This is likely an effect of the randomness observed in exome sequence variation, and the variation in HLA binding affinity of the resulting alloreactive peptides, and illustrates the potential for variability in alloreactive antigen presentation between different donors and recipients who undergo SCT.

**Figure 2.**
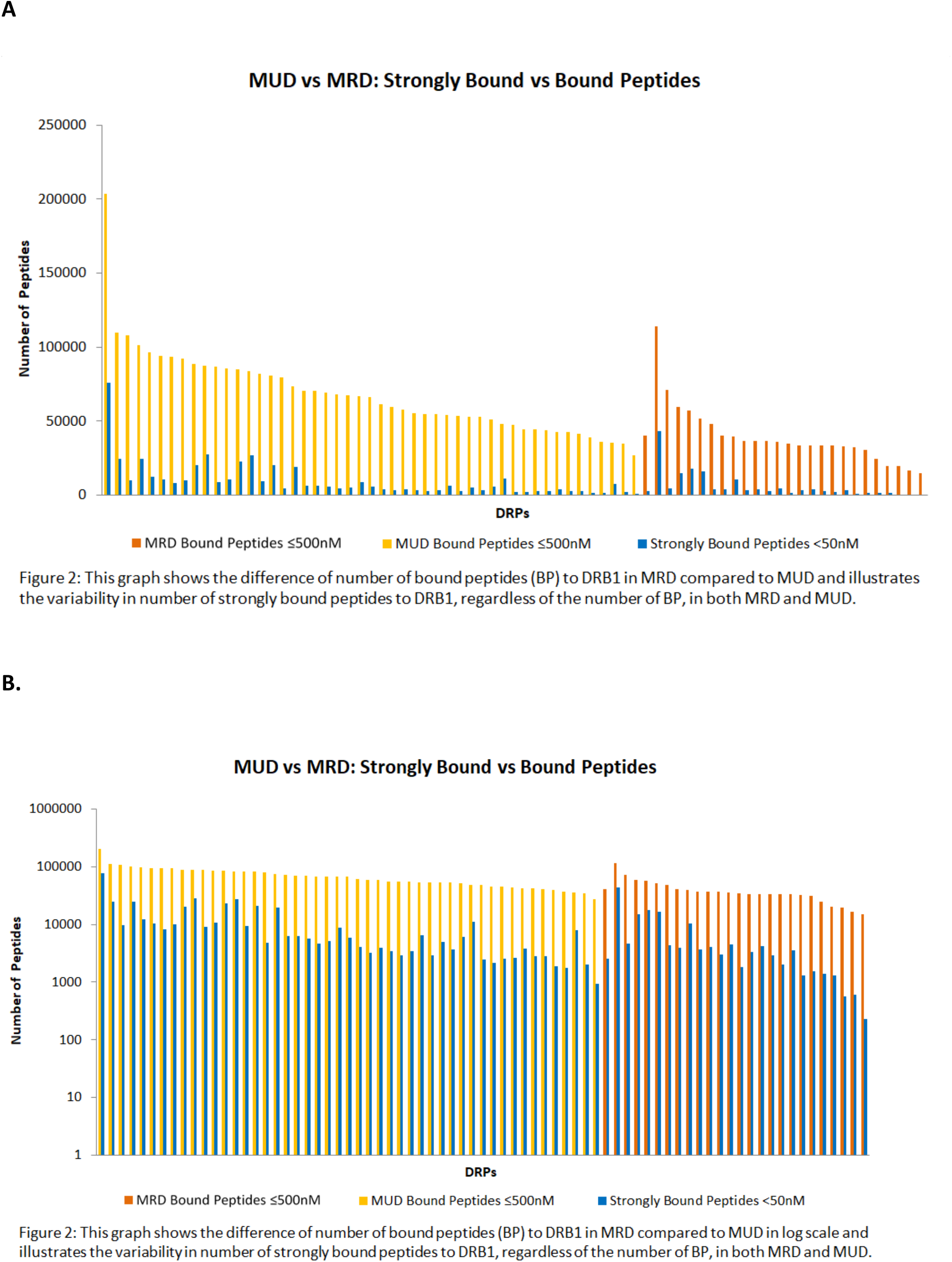
HLA class II bound peptides in HLA MRD and MUD DRP. Depicting SB and BP on standard (A) and logarithmic scales (B). Figure 2: This graph shows the difference of number of bound peptides (BP) to DRB1 in MRD compared to MUD and illustrates the variability in number of strongly bound peptides to DRB1, regardless of the number of BP, in both MRD and MUD. Figure 2: This graph shows the difference of number of bound peptides (BP) to DRB1 in MRD compared to MUD in log scale and illustrates the variability in number of strongly bound peptides to DRB1, regardless of the number of BP, in both MRD and MUD.

### Comparing HLA class I and II bound alloreactive peptides

The HLA class II binding peptides libraries were compared to previously-determined numbers of BP and SB on all Class I HLA alleles for the same patients. On average, the number of alloreactive peptides bound to the two HLA-DRB1 alleles with an IC50<500nM, was far greater than the number bound to the HLA class I loci (all 6 HLA-A, B & C alleles). Significantly more peptides bound HLA DRB1 molecules compared to all the HLA class I molecules put together; BP for HLA DRB1 median 52,983 compared with BP for all HLA class I molecules 4,532, yielding a median ratio BP-HLA class I/BP-HLA DRB1 per DRP of 0.09 (0.03-0.29; t test p<0.0001). The same trend was observed with SB with a median ratio of 0.23 per DRP (0.02-4.48; p =0.0001) (Figure 3A). There was correlation between the number of BP and SB for both HLA class I and to a lesser extent in HLA class II molecules in the DRP studied; Pearson correlation coefficient, R 0.71, p<0.0000001 for HLA DRB1 & 0.94, p <0.0000001 for all HLA class I molecules together (Figure 3B). Nevertheless, HLA class II molecules presented an order of magnitude greater number of peptides. There was little overlap in the binding affinities of various alloreactive peptides to different HLA class I molecules (Table 1B). The difference observed in HLA class I and II antigen presentation is likely a consequence of the larger peptide length presented on the dimeric HLA class II molecules. This increases the size of the peptide pool on offer (9 alloreactive peptides/SNP for HLA I vs. 15 for HLA II), and consequently the likelihood that alloreactive peptides will be presented. Tissue expression of the proteins from which the peptides presented on HLA class I, were derived was also determined and marked variation was observed in the RPKM values of the proteins of origin (Tables 1A & 1B).

**Figure 3.**
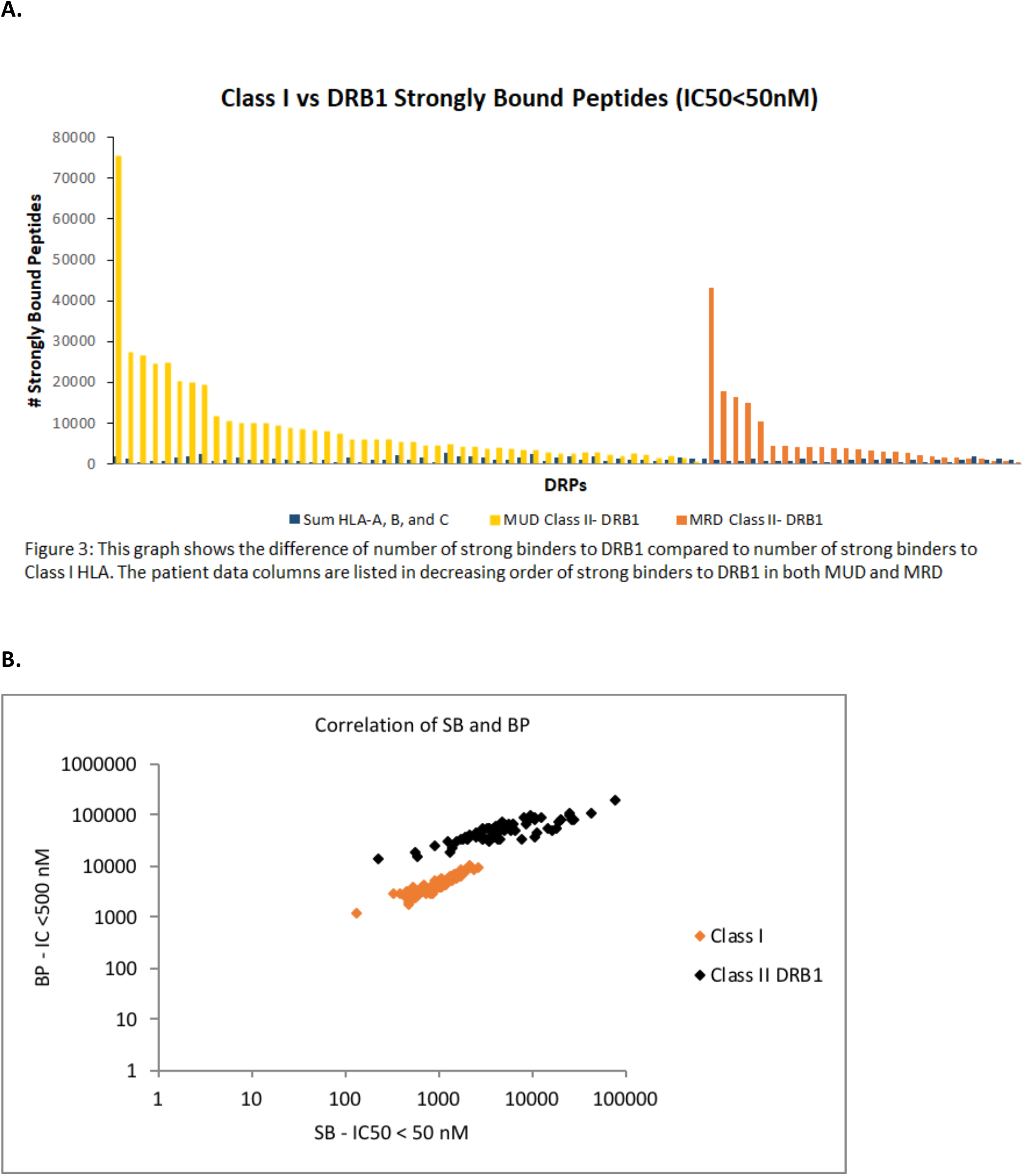
Comparison of HLA class I & class II bound peptides in HLA MRD and MUD DRP. Depicting SB on standard scale (A). Correlation of SB and BP for both HLA class I and class II molecules in MRD and MUD DRP. Figure 3: This graph shows the difference of number of strong binders to DRB1 compared to number of strong binders to Class I HLA. The patient data columns are listed in decreasing order of strong binders to DRB1 in both MUD and MRD

One DRP (# 26), was analyzed to determine the likelihood of peptide presentation from the same proteins on both HLA class I and II molecules. This would result in activation of both CD4+ and CD8+ T cells in the tissues expressing that protein, and greater potential for tissue injury. A comparison of strongly bound peptides (IC50 ≤50nM) demonstrates that this DRP had 143 genes, that yielded peptides binding both HLA class I and HLA class II. Different degrees of sequence of homology between these 9-mer and 15-mer peptides was observed (Table 2 & Supplementary Figure 1). This overlap suggests that if the degree of exome sequence variation in a DRP is sufficiently large, it is plausible that most tissues will potentially present mHA to both helper and cytotoxic T cells.

**Table 2.**
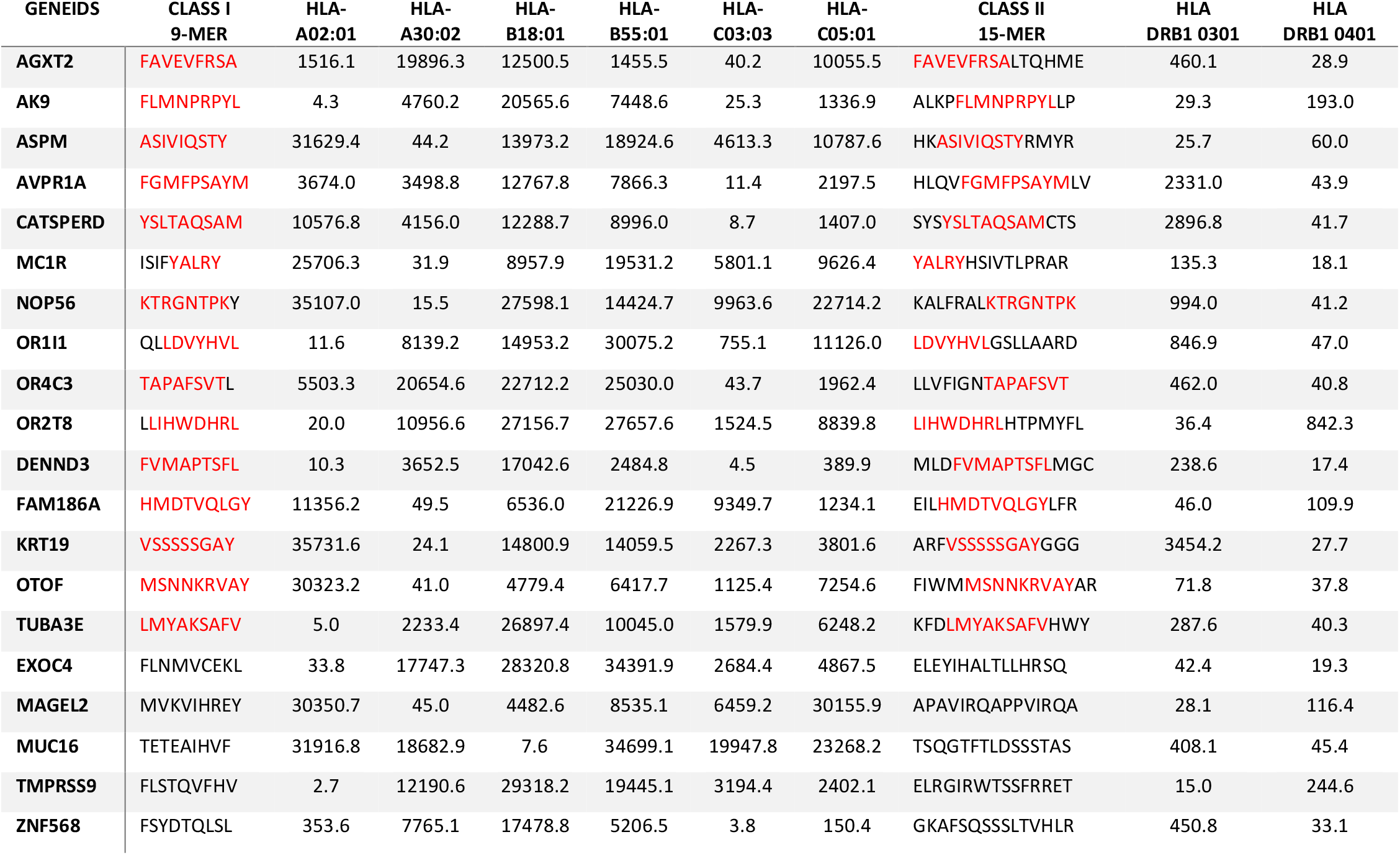
DRP 26, polymorphic HLA-bound (both class I & II) peptides derived from the same gene, with binding affinity values in the IC50, 0-50 nM range (highlighted in green). Varying degrees of sequence identity is seen between the HLA bound peptides; 5 peptides per degree of sequence homology are presented and homologous sequence is given in red. Rows 1-5, 9-mer HLA class I-bound peptide nested within 15-mer HLA class II-bound peptide; rows 6-10, sequence overlap between 9- and 15-mer peptides; rows 11-15, variable sequence homology (at least 1 nested, shown, and at least 1 matched only by gene nearby with no sequence identity, not shown); rows 16-20, different HLA bound peptides from the same gene with no sequence identity (Supplementary Figure 1).

### Demographic factors influencing HLA class II bound alloreactive peptides

Finally, demographic factors, including race and gender, that impact genetic disparity were analyzed. African-American vs. Caucasian DRP demonstrated a non-significant trend for increased HLA-DRB1 bound mHA in African American DRP, for both BP (74,179 vs. 53,735 in Caucasian patients; p=0.075) & SB peptides (11,972 vs. 7,503; p=0.36). There was no significant difference in the number of BP or SB in the gender-matched male or female DRP, not accounting for Y chromosome disparity in male patients receiving transplant from a female donor.

## Discussion

The data presented in this paper illustrate the large potential that HLA class I and especially HLA class II molecules have for recipient peptide antigen presentation in the context of allogeneic SCT. The magnitude of this antigen burden across the patient population makes it difficult to predict which patient will have a poor outcome. However, while in and of themselves these parameters may not yet be definitive for GVHD prediction, given the uniformly large magnitude of mHA identified in the patient cohort examined, these measures if appropriately analyzed may give insight into the quantitative principles of the T cell immune response. The following discussion gives a quantitative perspective of the impact the presentation of recipient antigens on HLA class I and II molecules may have on donor T cell responses following allogeneic SCT. The Dynamical System model of alloreactive T cell growth previously developed for HLA class I-presented-mHA is further developed for use in future studies to simulate universal alloreactive T cell responses.

An important clinical question in transplant immunology is how data from next generation sequencing (NGS) and novel machine-learning algorithms can be used to help identify optimal donors for SCT. To do this it is imperative to understand the quantitative principles at work in donor immune response and use these principles to develop methodology to simulate transplants with different donors *in silico*. Such simulations may then be used to identify both the ideal donor and the level of immunosuppression needed for optimal clinical outcomes. The mHA prediction methodology presented previously and extended herein, augmented by analysis of peptide cleavage sites to more accurately determine the probability of the generation of specific HLA binding alloreactive peptides may allow this prediction in the future^28^. As a first step towards this goal, it was shown that donor CD8+ T cell growth simulations may identify patients at risk for moderate to severe GVHD, however these associations were relatively weak^23^. While, one possible explanation for this is the stochastic nature of alloreactive antigen presentation on HLA molecules (both alloreactive and non-alloreactive peptides may bind HLA), an important limitation in the special case of the model described (HLA class I antigen presentation) was its lack of information on HLA class II mHA presentation and consequent CD4+ helper T cell responses in the donor-recipient pairs involved. Normally, CD4+ T helper cells play an important role in the homing of cytotoxic T cells to infected tissues, and in the case of GVHD to the target tissues^29, 30, 31^. In the transplant setting, T helper cells will recognize their target alloreactive antigens bound to HLA class II molecules; notably, these differ from the antigens recognized by CD8+ cytotoxic T cells and presented by HLA class I. The T helper cells initiate signaling by secretion of appropriate cytokines (IFN-γ, IL-2, IL-12, IL-17 etc.) and set up the homing signal for the cytotoxic T cells to invade the target tissue (Figure 4), which cause tissue injury through direct cytolytic activity. In the present study we estimate the magnitude of alloreactive antigen burden encountered by donor cytotoxic T cells and helper T cells in HLA matched DRP. This estimate may allow a more accurate calculation of the likelihood that a patient may develop T cell mediated tissue injury following SCT, then was previously possible^23^.

**Figure 4.**
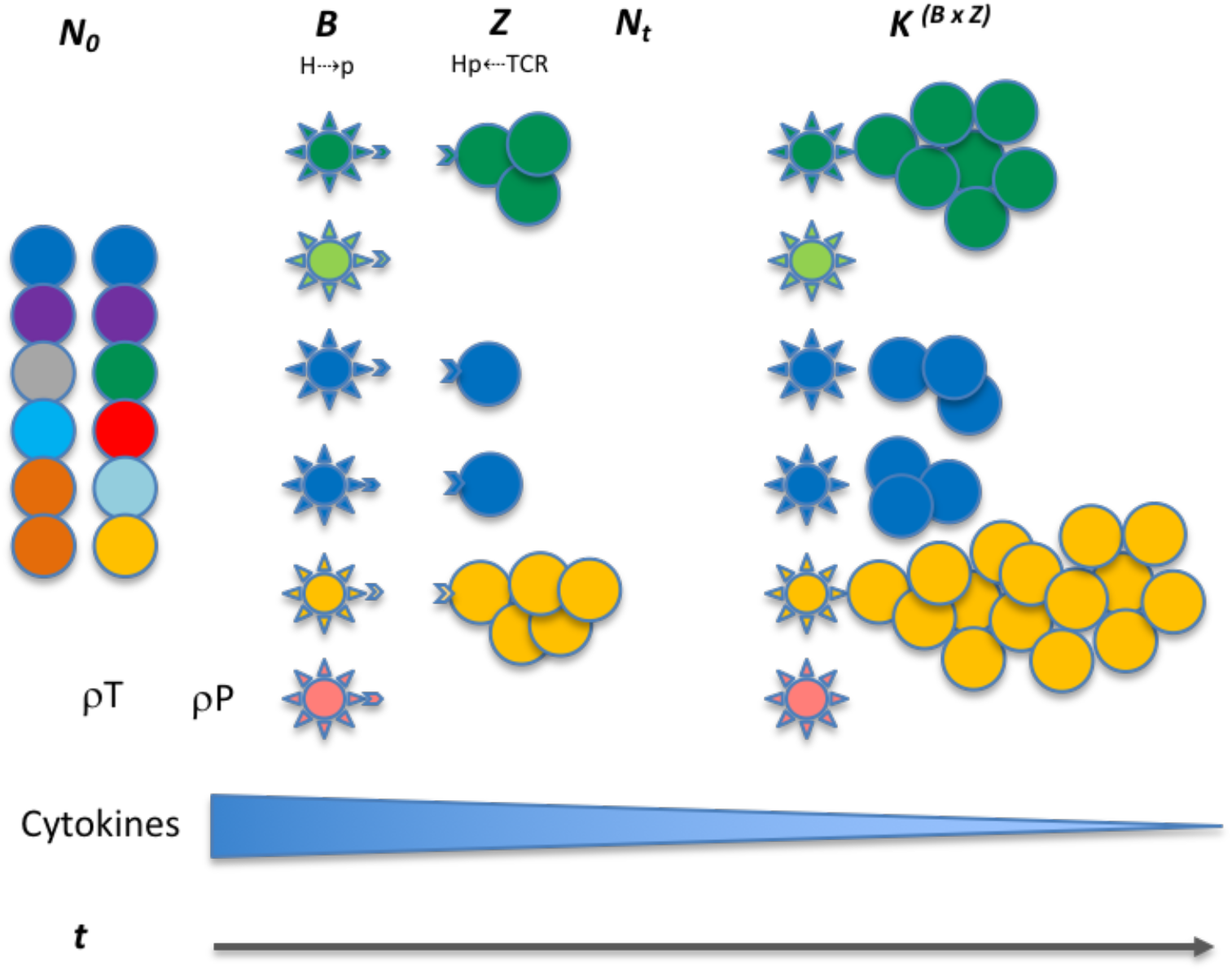
Interaction between donor T cells and recipient antigens presented on APC. Colored circles show different T cell clones with variable antigen affinity. The spiked spheres show antigen presenting cells with HLA molecules, colors indicate unique peptide antigens and correspond to T cell recognition. T cell growth is indicated over time in response to various influences, i.e. peptide affinity for HLA molecules (Vector *B*), and TCR affinity for HLA-peptide complex (vector *Z*), both leading to T cell growth. The infused allograft contains T cells with TCR of varying peptide antigen specificity (different colors), these may encounter peptides for which they have affinity (no color difference between APC and T cells), leading to growth. This variability makes antigen response a probability function of the likelihood of T cell presence and peptide antigen presentation (*ρP & ρT*). The expression level is depicted by more cells presenting the same antigen, making up for weak affinity driving proliferation. The cytokines made by APCs get taken up by the cytokine receptors on the T cells, leading to a diminishing effect as the T cell repertoire expands.

### T cell clonal proliferation in response to mHA-HLA complexes: The logistic equation of growth

Previous work has shown there to be far greater diversity in the T cell repertoire of CD4+ T cells than in the CD8+ T cells in the post-transplant period in both allogeneic and autologous SCT^32^. In fact, CD4+ T cell diversity has been found to be about 50 times greater than CD8+ T cell diversity^33^. The relative magnitude of antigen presentation by HLA class II compared with HLA class I molecules allows one to understand this difference in clonal diversity between the helper and cytotoxic T cells. The ability of HLA class II molecules to present larger peptide sequences is related to their structure compared to HLA class I molecules. The antigen-binding region of HLA Class II molecules consists of both an invariant α and a variable β domain, whereas that of HLA Class I molecules contains only α domains resulting in the binding of a wider range of peptide sequences^6 34^. This differential antigen presentation results in the quantitative difference observed between the two classes of T cells and may be understood using the dynamical systems approach. In this model, growth equations have been used to simulate the cytotoxic T cell growth in response to HLA class I presented antigen,

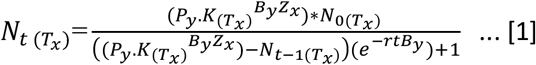

This iterating equation describes the logistic growth of a CD8+ T cell clone *T_x_* in a polyclonal T cell graft infused into a recipient (Figure 4 & Supplementary Table 2). *N*_*0* (*Tx*)_ is the T cell count at the time of transplantation (assumed to be 1 for this equation), *N*_*t* (*Tx*)_ is the T cell count after *t* iterations (time) following SCT. *N*_*t-1* (*Tx*)_ represents the T cell count for the previous iteration and *K* is the constant that will determine the T cell count at the asymptote (steady state conditions after infinite iterations), *K* _(*Tx*)_, representing the maximum T cell count the *system* would support (*carrying capacity*); *r* is the growth rate. In the logistic equation, the steady state count for each T cell clone (*K^BZ^*) will be proportional to the <u>product</u> of the binding affinity of the target peptide mHA (peptide *y*) for the HLA molecule (*afmHA* = 1/IC50 in Koparde *et al*, in this paper, *B_y_* for peptide *y*) and the affinity of T cell clone, 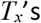 T cell receptor for the mHA-HLA complex (*afTCR* = *1/IC50* in Koparde *et al*, now *Z_x_* for T cell clone *Tx*)^23^. In this model, the parameter *r*, determines the growth rate of the specific clone and reflects the effect of the costimulatory molecules and cytokines driving T cell proliferation. This iterating equation gives instantaneous T cell count (magnitude of the proliferative response) in response to antigens presented. The tissue expression of proteins from which peptide *y* is derived (*P_y_*) is a coefficient/multiplier for the steady state T cell population *K^BZ^*, and may be estimated by RNA sequencing techniques, and reported as Reads or Fragments Per Kilobase of transcript per Million mapped reads (RPKM or FPKM)^35^. In real-world situations the term *P_y_* will have a time modifier, *e^t^*, associated with it, as protein expression and antigen amount declines over time because of tissue injury. This time relationship will be ignored for simplicity at this time. It is important to recognize that in HLA class I-presented antigen-driven T cell expansion, this term is utilized in its entirety given that HLA class I molecules are loaded using peptides derived from proteins present in the cytosol. This however is not the case for HLA class II molecules, which present antigens endocytosed from the extracellular environment^36^. This means that when calculating helper T cell growth, the term *P* will be modified to *P.c*, with a constant, *c*, reflecting the attenuation of antigen concentration given its ‘scavenged’ nature as opposed to direct cytosolic presence, in other words, *0* < *c* < *1* (for CD8+ T cells, *c*=*1*). Thus, the equation for determining helper T cell growth will take the general form,

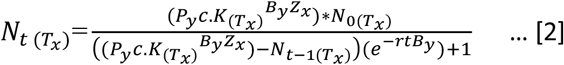

Adjusting the variable *P* means that the absolute magnitude of the steady state T cell population for each of the dominant (high-ranked) helper T cell clones will be smaller than that for each of the dominant cytotoxic T cell clones, nevertheless because of the greater number of antigens presented by HLA class II molecules there will be a greater number of CD4+ T cell clones, and thus greater clonal diversity of helper T cells when compared to cytotoxic T cells. This also means that in a Power law clonal frequency distribution analysis^37, 38^, the contribution of the highest-ranking (most numerous) T cell clones to the entire repertoire will be higher with cytotoxic T cells. Conversely, in the T helper cell population there will be a larger number of high-ranking clones which contribute a larger component of the overall repertoire. Given the greater number of antigens there may be greater competition between the clones, which in a model accounting for competition between clones will lead to slower growth of helper T cell clones, a relatively frequent clinical observation^39^. Also, given the restriction of HLA class II molecules to antigen presenting cells the absolute magnitude of steady-state helper T cell clonal populations will be smaller; however, since HLA class I molecules are expressed on all nucleated cells, cytotoxic T cells get a proliferative signal from many different cell types, therefore steady state T cell clonal counts can be further augmented. From an evolutionary and T cell response sensitivity and specificity standpoint, it is logical that the cytotoxic T cell-recruiting signal provided by CD4+ T helper cells should be more sensitive, triggered by a greater variety of antigens, but when it comes to actual tissue destruction by CD8+ cytotoxic T cells, a more fine-tuned HLA class I bound, shorter peptide with greater specificity required for presentation, provides the necessary stimulus. This would come from the prevention of non-specific binding of peptide antigens to the more ‘discriminating’ HLA class I molecules.

### Quantifying mHA-HLA-TCR interactions: On matrices, vectors & tensors

Following the above general discussion about T cell behavior, it is necessary to develop a model that will account for the potentially large arrays of antigens being presented in allogeneic SCT. As noted earlier, immunotherapy and SCT are fraught with the risk of treatment failure either in the form of relapsed malignancy or immune mediated normal tissue injury (GVHD or graft rejection). Various outcome prediction algorithms and models have been developed using increasingly sophisticated characteristics studied statistically^40, 41^. These may allow improvement in clinical outcomes prediction, but often do not provide mechanistic insight into the reason for the observed clinical outcomes. Further, while principles of immune therapy and the mechanisms of T cell action are well known from work on mouse models and *in vitro*^42, 43^, when the antigenic complexity encountered *in vivo* in human SCT recipients is considered, the existing models do not reliably predict individual clinical outcomes. This is also true of the T cell repertoire that emerges following SCT.

Nevertheless, mathematical methods are available that have long been used in physics to understand natural phenomenon and may be extrapolated to biological systems such as immune response modeling. For example, the concept of vectors and operators has been used to simulate aggregate T cell clonal responses to antigen arrays^22, 23^. However, this method is limited in that it requires identification of unique mHA-HLA and cognate TCR for application. To overcome this limitation, a related mathematical method, tensor analysis, may be used to simulate the immune responses to the vast library of tissue specific antigens presented by the entire spectrum of HLA molecules in an individual. In physics, tensors describe interaction between vector quantities and their components, so they enable determination of variation in vector magnitude and direction and subsequent mapping to a different ‘state’. In other words, tensors help describe vector transformation when multiple forces are acting upon an object, which itself may be a vector^44, 45, 46^. It is important to recognize that these methods have been developed for use in ‘linear’ physical systems, however biological systems are seldom linear. They follow nonlinear dynamics such as Power laws and exponential growth patterns, which require development of methodology which can account for the complexity in biologic systems because of the multiplicity of variables encountered. It is for this reason that tensor methodology may lend itself to the study of the alloreactive immune response problem. In the example at hand, the donor T cell array infused into the recipient may be considered as a vector, which is modified by the interaction between the T cell receptors (TCR) on the donor T cell clones and the recipient mHA-HLA complexes and is transformed to a new state following SCT. The interacting TCR & mHA-HLA complex in this example may be considered as a tensor, modifying the T cell clonal vector. Tensors remain invariant in different frames of reference and in this application of the concept, the mHA-HLA-TCR interactions, determined by the protein sequences remain constant, regardless of tissues and individuals where the interactions may be occurring. In other words, the unique peptide sequences’ affinity to specific HLA molecules and TCR will remain the same across individuals and tissues. In essence, such an alloreactivity tensor comprised of recipient mHA and HLA, in the presence of donor T cell repertoire influences the relative growth of alloreactive T cell clones versus the non-alloreactive clones. Accordingly, clinical GVHD may or may not manifest.

To understand this notion, consider a basic adaptive immune response to a recipient alloreactive peptide following SCT (or any other antigenic peptide); the first interaction is between the alloreactive recipient peptide and the HLA molecule resulting in the binding and presentation of the peptide on the HLA molecules (Figure 5). Consider two HLA molecules *H*_*1*_ and *H*_*2*_, and two peptides *p*_*1*_ and *p*_*2*_, each recognized by only one of these two HLA molecules; a matrix may be constructed showing the peptides bound to the relevant HLA molecules^47^. The possible interactions between the peptides *p*_*1*_ and *p*_*2*_ in a system of two HLA molecules *H*_*1*_ and *H*_*2*_, may be depicted in matrix form as follows.

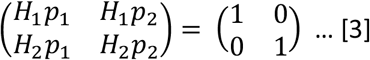

**Figure 5.**
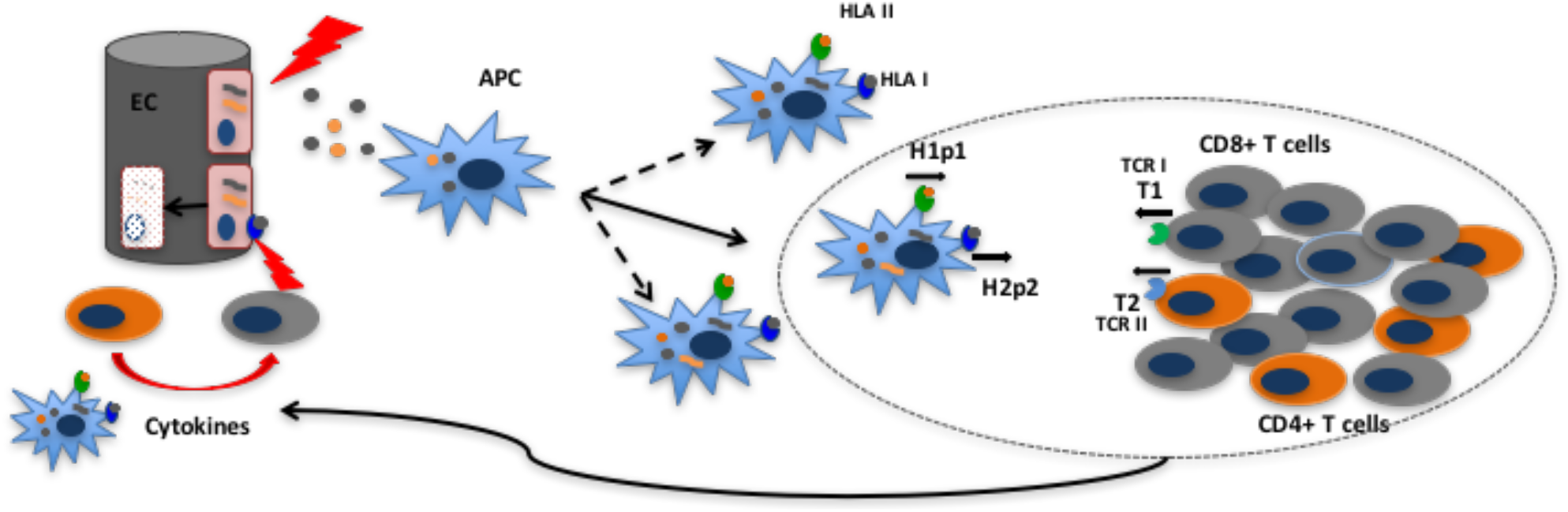
Tissue injury releases polymorphic recipient mHA from epithelial cells (EC); these and endogenous antigens are presented by APC; the APC proliferate and migrate to the lymph node triggering a CD4+ and CD8+ T cell clonal expansion according to the logistic equation of growth. These T cell clones then enter the circulation and migrate to the tissues to initiate tissue injury. Short black arrows in the oval (lymph node) below H1P1 and TCR1 indicate affinity vectors B and Z respectively.

**Figure 6.**
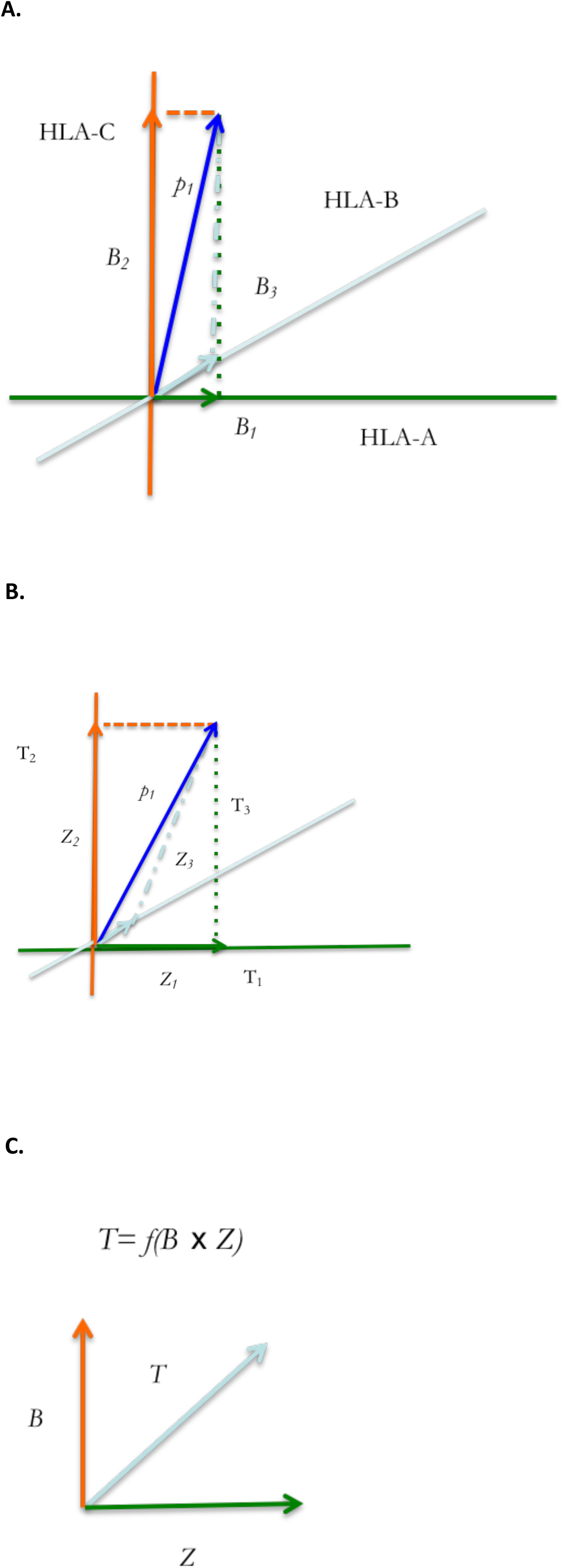
Tensor diagram for the HpT tensor. Peptide *p_1_* binds different HLA molecules, A, B and C with affinities, B_1_, B_2_ and B_3_ (A), while each TCR (TCR_1-3_) binds these HLA-peptide complexes (with peptide, *p_1_*), with affinities Z_1_, Z_2_, and Z_3_ (B). Blue arrows indicate average effect of multiple binding affinity vectors. These affinities will remain unaffected in different tissues and individuals and T cell clonal growth in response to the polymorphic peptide is proportional to their product (C).

The 0 and 1 represent conditionality of interaction between the peptides and HLA. The matrix on the left-hand side of equation 3 represents vector quantities, *H_1_p_1_, H_1_p_2_, H_2_p_1_ or H_2_p_2_*, which have a magnitude (binding affinity, expressed in 1/IC50, nM^-1^) and a ‘direction’ given by the specificity, i.e. unique affinity of the peptide for the HLA molecule. Given affinity of H_1_ for p_1_ and H_2_ for p_2_, this interaction yields an identity matrix. The interaction between the peptides and HLA molecules constitute a matrix where peptide recognition and binding by an HLA molecule is represented by 1, and the converse situation by 0. Thus, the numbers 1 & 0 represent the selectivity of peptides with a certain sequence (and commensurate length) for specific HLA and *vice versa*. These two alloreactive HLA-peptide complexes may then be presented to donor T cell clones by the antigen presenting cells, (Figure 4) and specific donor T cell receptors may recognize these unique HLA-peptide combinations and bind. In this example, *TCR_1_* only recognizes *H_1_p_1_* and, *TCR*_*2*_ only recognizes *H_2_p_2_*. The resulting matrices are given below

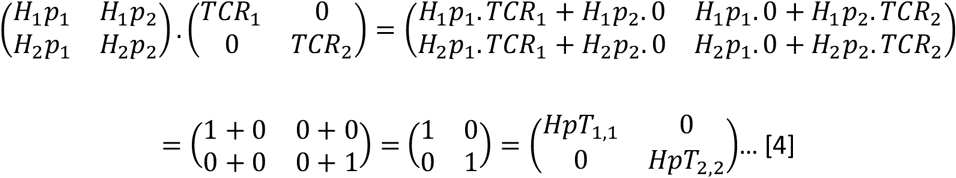

The right-hand side of equation 4 is a tensor with two vector quantities, the affinity of HLA molecule for the peptide and the affinity of the TCR for the peptide-HLA complex, which may be summarized as follows

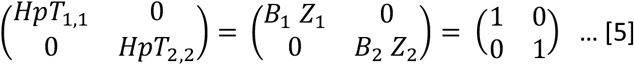

The matrix depicted in equation 5, is a tensor of the second rank with two vector quantities, i.e. the affinities *B* and *Z* (specific binding between HLA & peptide (*B*) and between HLA-peptide & TCR (*Z*)), which are depicted by *HpT_1,1_* and *HpT_2,2_. HpT* in this case symbolizes the HLA molecules, peptides and TCR interacting with each other, and the subscripts *1* and *2* are called indices in tensor terminology, identifying interactions between specific molecules (e.g., *p_1_* and *p_2_*). The identity matrix reflects the affinity of specific TCR for specific mHA-HLA combinations. It is to be noted that, the same peptides given above may bind other HLA molecules with a different affinity and there may be TCR which bind these alternative antigen complexes with different affinities, constituting different vectors (Figure 5A & 5B). Along the same lines, a given peptide or TCR may interact with different partners yielding different *vector components*. For example, in the above matrices, *TCR_1_* may interact with both *H_1_p_1_* and *H_1_p_2_*, the magnitude of the former will be 1 and the latter, 0. However, given the continuous nature of the IC50s observed for different peptides with different HLA molecules in the analysis presented in this paper it is unlikely that the vector magnitudes are going to be binary in nature. The well-known phenomenon of immune cross reactivity is an example of the vector components which are not binary^48^. It is also important to note that the forces (vectors) represented by *B* (*H_1_p_1_*) and *Z* (*TCR_1_*) may be considered orthogonal (perpendicular) because their direction is imparted by the unique recognition of peptide sequence by HLA, and that of peptide-HLA complex by TCR respectively. Thus, the growth of the T cell clone resulting from this interaction may be considered a ‘cross’ product of these two forces (*Sin 90 °=1*, for orthogonal vectors) (Figure 5C).

### T cell vector transformation: Enter Operators

In the SCT context the alloreactivity tensor, *HpT*, determines the magnitude (and direction) of T cell clonal growth vector in response to antigens. T cell clones with receptors *TCR_1_* and *TCR_2_* respectively will grow in response to the *HpT* Tensor. It is to be noted that the HLA-peptide driven T cell clonal growth vector is distinct from the TCR affinity vector for HLA-peptide complex, even if one considers that mHA-HLA affinity vector drives T cell clonal growth of the relevant TCR bearing clone. This relationship is analogous to applied force, resulting in motion at a certain velocity and consequent mass displacement which are distinct vector quantities pointing in the same direction (with time being the scalar distinguishing between them; T cell clonal growth is also a time-dependent function). In the above example, the T cell clonal growth vectors, comprising the two T cell clones bearing the T cell receptors *TCR_1_* and *TCR_2_*, are termed *T_1_* and *T_2_* respectively. These constitute a vector matrix, which is transformed over time *t* by the *HpT* tensor to the vectors 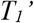 and 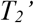.

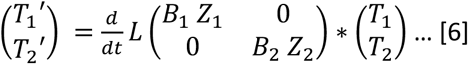

In equation 6, the vector 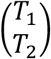 *is* transformed by the *HpT* tensor and the logistic operator, 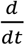 *L* previously defined as the logistic equation for T cell growth, which incorporates the term *B_y_Z_x_* included in the *HpT* tensor,

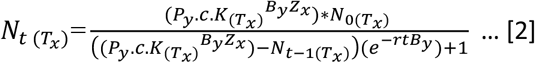

### T cell growth: the effect of co-stimulation, checkpoints and cytokines

In equation 2 the term *r* quantifying growth rate is an aggregate measure of different influences on T cells and may be considered a scalar multiple of a tensor quantity. This term represents the cumulative growth effect of the costimulatory and inhibitory molecules present on the T cells and the cytokines present in the environment. In the dynamical system model of T cell growth, the T cell steady state numbers are determined by TCR-mHA-HLA affinity (*BZ*), also called ‘Signal 1’. A second critical influence on T cell growth is provided by ‘Signal 2’ mediated by the costimulatory molecule CD28 and inhibitory molecule CTLA4 (*S2*) may be mathematically represented by, CD28 = 1, CTLA4 = 0. Additionally, the checkpoint mechanism (*CP*) comprising the PD1 receptors, if engaged may be represented by a variable valued at 0 because no T cell growth will occur, and when absent, valued at 1. Finally, ‘Signal 3’, (*S3*) represents the effect of cytokines on T cell growth (Supplementary Figure 2)^49, 50, 51^. Considering that all these variables contribute to T cell growth, the term *r* is therefore a composite of the following factors,

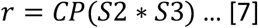

Solving this equation for lack of PD1 engagement (1) and the presence of CD28 expression (1) yields,

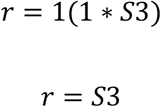

Solving the equation for CTLA4 expression or PD1 engagement gives *r* a value of 0, which yields *e°* = 1 in equations 1 & 2, consistent with suppression of T cell growth. In other words, the presence of PD1 engagement by PDL-1 or the engagement of CTLA-4 instead of CD28, by CD80 on APC, changes *r* to zero, eliminating the effect of time *t*, which changes the value of *e* to 1 (in equation 2), leading to growth arrest of the T cell clone.

As for *S3*, the cytokine mediated signal may also be considered a second order tensor quantity, consisting of a matrix with cytokines and cytokine receptor vectors, because the cytokines and their receptors, have different magnitudes and varying receptor specific effects (directionality) on T cell growth and differentiation. Ignoring the di- or trimerization of cytokine-receptor protein subunits, a simplified version of the cytokine tensor may be constructed as follows,

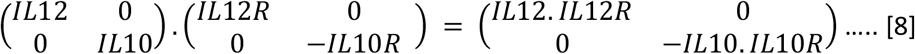

This is the cytokine tensor, *Ck*, with the example showing the interaction between IL-12 and IL-10 and their respective receptors. It should be noted that cytokines may bind related receptors with different affinities, providing different vector components. The negative sign means a growth suppressive effect, the net effect of cytokines can either be negative or positive and as a multiple of the CD28-PD1 expression term, the *Ck* can alter the magnitude and direction of effect of the exponent in equation 2 (by changing the symbol of *r* from - to +). Equation 7 therefore is modified to

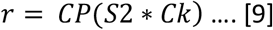

Further complicating these estimations from a physical standpoint at a cellular level in equation 8, cytokine exposure will be variable since these effects are ‘local’ to the tissue or lymph nodes. Cytokines likely depend on diffusion via capillary action in the extracellular matrix to create a ‘field’ in which the T cells experience the cytokine effects. These effects on growth are of an exponential nature because of *r* being an exponent in equations 1 & 2^52^. The receptor expression levels also vary on different cells and confer a direction by means of influencing differentiation and functional specificity to the T cell clones with unique TCR.

### Evolution of the T cell repertoire: Putting it all together

The above discussion illustrates the complexity inherent in the multiple factors influencing the T cell responses to antigens presented by HLA molecules. Nevertheless, it makes it clear that despite the complexity, it is possible to describe the immune interactions in mathematical terms, and therefore it is also possible to simulate them, especially when antigen presentation data are available. To do so one may take the example of a random collection of tissue associated peptides. First, consider an alloreactive peptide of any size varying between 7-18 amino acids. This peptide will have a choice of binding to HLA class I and II molecules (there are six of each). Therefore, depending on its size and mode of acquisition (extracellular or cytosolic) it will bind to the relevant HLA molecules with a unique binding affinity. It is to be noted that depending on the number of binding HLA molecules and the concentration of competing peptides, there will be a probability function associated with each of these interactions. As demonstrated above in equations 4 and 5, the mHA (polymorphic peptide) binding affinity to available HLA molecules, may be considered to represent the components of the immune response vector to this antigen (or degrees of freedom for the peptide). For most peptides, only one component (one HLA-mHA complex) with the strong interaction will be relevant, and others with weak interactions may be ignored. With the peptide bound to one of the HLA molecules (or more depending on binding affinity with other HLA molecules), it is presented on the APC. If a T cell clone with a TCR which has affinity for the HLA-peptide complex is present (a second probability term), then depending on the CD28/CTLA-4 and PD1 expression levels in the T cell clone, it will grow in the cytokine ‘field’ present in the tissue.

Thus, consider peptides (*p_1_, p_2_*…*p_n_*) with high affinities *B_1_, B_2_*…*B_n_* for HLA molecules *H_1_, H_2_*…*H_n_* respectively, but with a very low-level affinity for the non-corresponding HLA molecules present in the individual (e.g., the components *p_1_H_2_, p_2_H_n_, p_n_H_1_*, not considered here for the sake of simplicity in illustration, but fundamental to the tensor concept). These mHA-HLA complexes have corresponding T cell receptors *TCR_1_, TCR_2_…TCR_m_* with affinities, *Z_1_, Z_2_…Z_m_*, the tensor *HpT* may be written as follows,

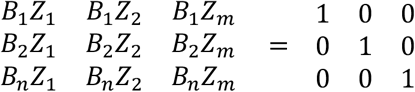

Here *n* and *m* are indices which indicate the HLA-peptide affinity (*Bi*) and TCR binding affinity to the HLA-peptide complex (*Zj*). This is the *alloreactivity tensor*, and it reflects the interaction of the alloreactive peptides with the HLA molecules in that individual and transforms the T cell clonal vector comprised of the array of the T cell clones bearing the above TCR <*T_m_*> according to the logistic function.

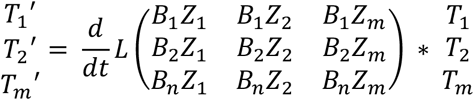

This results in the transformation of the infused donor T cell repertoire, with *T_1_, T*_*2*_…*T_m_* being transformed to 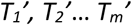 following transplant. The logistic growth equation provides the rule for transformation, so equation 1 may also be rewritten as follows for the *i^th^* HLA-bound-peptide, *p_i_*, and the responding *j^th^* T cell clone <*T_j_*> in a repertoire comprised of T cell clones *T*_*1*_ thru *T_m_*.

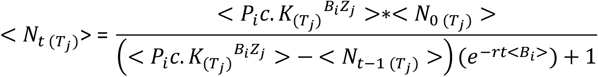

Substituting the value of *r* from equation 9 in this equation, we get,

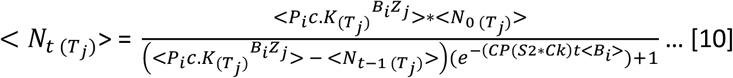

The aggregate alloreactive T cell response at time, *t* is then

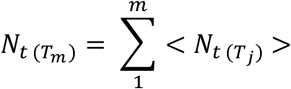

This general equation describes the transforming effect of the alloreactivity tensor and the cytokine tensor on the T cell repertoire following SCT. The risk of alloreactivity developing clinically will in this instance be proportional to *N_t(TM)_*.

### Dynamical system model of alloreactive T cell response and clinical observations

Does this model explain observations in clinical transplantation? To determine this one may consider the matter of HLA-DPB1 mismatching and alloreactivity in 10/10 HLA matched DRP^53, 54^, and for that matter the general problem of HLA mismatched SCT and associated negative clinical outcomes^55^. In the dynamical system model this phenomenon may be easily understood; the mismatched HLA DPB1 epitopes are highly expressed so instead of having a fraction of the protein expressed (the term *P.c* in Eq. 2) governing CD4+T cell clonal growth, T cell clones bearing TCR that recognize epitopes on HLA DPB1 encounter an order of magnitude higher target concentration with a marked amplification of the steady state alloreactive T cell clonal populations compared to a standard HLA class II bound mHA. Indeed, polymorphisms impacting the level of HLA DPB1 expression correlate with the likelihood of GVHD developing^56^. Further any peptides bound to the mismatched HLA will be novel antigen complexes for the donor T cell clones to recognize. This would result in a strong aggregate immune response to the mismatched HLA (and its presented peptides) which is widely expressed, and this response is significantly larger than a mHA-HLA directed immune response.

Despite the ability of the model to explain some common clinical observations (logistic growth of T cells, power law distributions, and CD4/CD clonal distribution), it will not be validated unless it explains the random occurrence of GVHD following allografting. A discussion of this has previously been presented, (Koparde et al 2017) where the competition between non-alloreactive and alloreactive peptides for HLA binding and presentation was invoked as a possible reason for this, resulting in a probability distribution (*ρHp_n_*) for the alloreactive peptide *p_n_* to be presented on HLA molecule *H*. A further consideration in the development of GVHD from these alloreactive T cell clonal growth simulations is the probability function introduced by peptide cleavage potential, which affects the likelihood of antigen presentation, as well as whether the relevant T cell clones are present following transplantation (*ρT_m_*). The probability of peptide cleavage 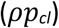 is determined by the amino-acid sequence at the C terminal of the peptide antigens^57^, as such, several peptides in our study may have low likelihood of presentation and may be ignored to simplify the model. The likelihood of alloreactive antigen response (*ρ_GVHD_*) may then be calculated as

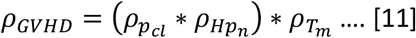

Computed for each alloreactive peptide, the probability of clonal expansion of the mHA-targeting-T cells will be significantly diminished as the number of probability terms are introduced into the computations, which explains why despite many potential alloreactive antigens being present in each donor and recipient not every patient develops GVHD.

Another clinical phenomenon, the T cell growth amplification effect of cytokines is well recognized clinically. This is recognized in both the need for lymphodepletion prior to adaptive immunotherapy and in the cytokine release syndrome seen following it^58, 59^. Thus far in the dynamical system model discussed above the cytokine tensor effect has been described as modulating rate of T cell clonal growth. However, cytokines effect not only the rate, but they also effect the magnitude of clonal expansion, amplifying the T cell clonal growth. This may be modelled using the iterating equation

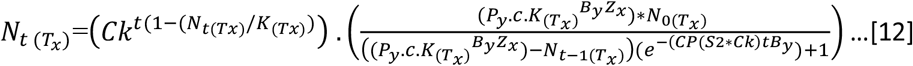

This equation demonstrates the effect of the cytokine tensor, *Ck*, as a time-dependent function, which in the beginning increases the magnitude of T cell clonal growth for clones expressing the relevant cytokine receptors by an order of magnitude. As the number of T cells increases, this time-dependent effect declines to a steady state level since the cytokines are taken up and utilized by the growing T cell population. This relationship plotted over time demonstrates the familiar T cell antigen response curve and mirrors the effect of antigen presenting cell growth previously described (Supplementary Figure 3) (See Koparde *et al*^23^, for discussion of APC-T cell interactions).

A final consideration in building this model is that the antigen matrices presented above are ‘identity matrices’ with binary values of, 1 along the diagonal of a square matrix and 0 elsewhere. In physiologic conditions, however there will be a continuum of values because of differential binding of peptides to various HLA molecules and cross reactivity of T cell receptors with such antigen complexes, generating *random number matrices*, rather than identity matrices^60^. This will add another element of complexity to the antigen-effector interactions, and possibly provides a rationale for complex GVHD phenotypes observed.

In conclusion, there is considerable genetic variation present between HLA matched transplant donors and recipients. *In silico*, this yields a putative large array of recipient mHA bound to both HLA class I and class II molecules, which when viewed from the frame of reference of responding donor T cells may be used to develop a mathematical model to allow simulation of the generalized T cell responses in allograft recipients. The quantitative understanding of alloreactivity thus gained may allow greater precision in donor selection and management of immunosuppression.

## Acknowledgements

This study conducted at Virginia Commonwealth University’s Massey Cancer Center was supported, in part, by research funding from the NIH-NCI Cancer Center Support Grant (P30-CA016059; PI: Gordon Ginder, MD) and by research funding from Virginia’s Commonwealth Health Research Board Grant #236-11-13 (PI: Michael Neale, PhD). We acknowledge Allison Scalora and David J Kobulnicky for collection of clinical outcomes data and analysis. Sequencing and Bioinformatics Analysis was performed in the Genomics Core of the Nucleic Acids Research Facilities at VCU, supervised by GB. VK performed bioinformatic analysis of the sequencing data to identify unique peptides and their HLA binding affinity, as well as tissue expression and wrote the paper. MS performed sequencing on samples identified and procured by CR. AS analyzed the data, performed statistical analysis and wrote the paper. AS, CH and MJ-L created data files with unique peptides and HLA IC50 values and wrote the paper. AT, MN, GB developed the WES study. AT developed the vector-operator and tensor models and wrote the manuscript. JR, RQ, MN, SH, DW and BAR critically reviewed the manuscript and provided expert commentary. All the authors contributed to writing the manuscript.

---

**Supplementary Table 1.** Demographics

| **Table 1. Patient Characteristics, n (%)** |  |  |
| --- | --- | --- |
| **Total Transplants** |  | **77** |
| **Gender** |  |  |
| **Male/Female** |  | **43 (55.8)/34 (44.2)** |
| **Patient Age** |  |  |
| **Median (range)** |  | **55.6 (21-73)** |
| **Donor** | **MRD** | **26 (34)** |
|  | **MUD** | **41 (53)** |
|  | **MMRD** | **1 (1)** |
|  | **MMUD** | **9 (12)** |
| **Stem Cell Source** | **Bone Marrow** | **7 (9)** |
|  | **Peripheral Blood** | **70 (91)** |
| **Diagnosis** | **Acute lymphoid leukemia** | **5 (6)** |
|  | **Acute myeloid leukemia** | **29 (38)** |
|  | **Chronic lymphocytic leukemia and lymphomas** | **15 (19)** |
|  | **Multiple myeloma** | **4 (5)** |
|  | **Myelodysplastic syndromes** | **24 (31)** |
| **Conditioning** | **Reduced Intensity** | **46 (60)** |
|  | **ATG/TBI** | **19 (25)** |
|  | **Busulfan/Fludarabine** | **27 (35)** |
|  | **Fludarabine/Melphalan** | **2 (77)** |
|  | **Myeloablative** | **31 (40)** |
|  | **Busulfan/Cyclophosphamide** | **17 (22)** |
|  | **Cyclophosphamide/TBI** | **11 (14)** |
|  | **Etoposide/TBI** | **1 (1)** |
| **GVHD prophylaxis** | **Anti-thymocyte globulin (ATG)** | **62 (81)** |
|  | **Cyclosporin A/Methotrexate** | **16 (21)** |
|  | **Cyclosporin A/MMF** | **4 (5)** |
|  | **Tacrolimus/Methotrexate** | **34 (44)** |
|  | **Tacrolimus/MMF** | **23 (30)** |
| **Race** | **Caucasian** | **60 (78)** |
|  | **African American** | **17 (22)** |

**Supplementary Table 2.** Explanation of variables used in the equations.

| Variable | Description | Previously |
| --- | --- | --- |
| $T$ | T cell clone | |
| $N$ | T cell count at various times (time indicated by subscripts) | |
| $K$ | Constant determining steady state T cell count | |
| $t$ | time/iterations | |
| $r$ | Growth rate | |
| $e$ | 2.7182, base of natural logarithms | |
| $p$ | Alloreactive peptide | |
| $H$ | HLA molecules | |
| $TCR$ | T cell receptor of T cell clone, $T$ | |
| $B$ | Binding affinity of peptide, $p$ for HLA molecule, $H$ | $afmHA^{23}$ |
| $Z$ | Binding affinity TCR for mHA-HLA complex ($Hp$) | $afTCR^{23}$ |
| $P$ | Tissue expression of protein with alloreactive peptide $p$ | $P_{exp}^{23}$ |
| $c$ | Constant which reduces $P$ for HLA class II presentation | |
| $HpT$ | Alloreactivity tensor | |
| $Ck$ | Cytokine tensor | |
| $CP$ | Checkpoint molecule | |
| $S2$ | Signal 2 | |
| $L$ | Logistic operator | |
| $\rho$ | Probability | |

**Supplemental Figure 1.**
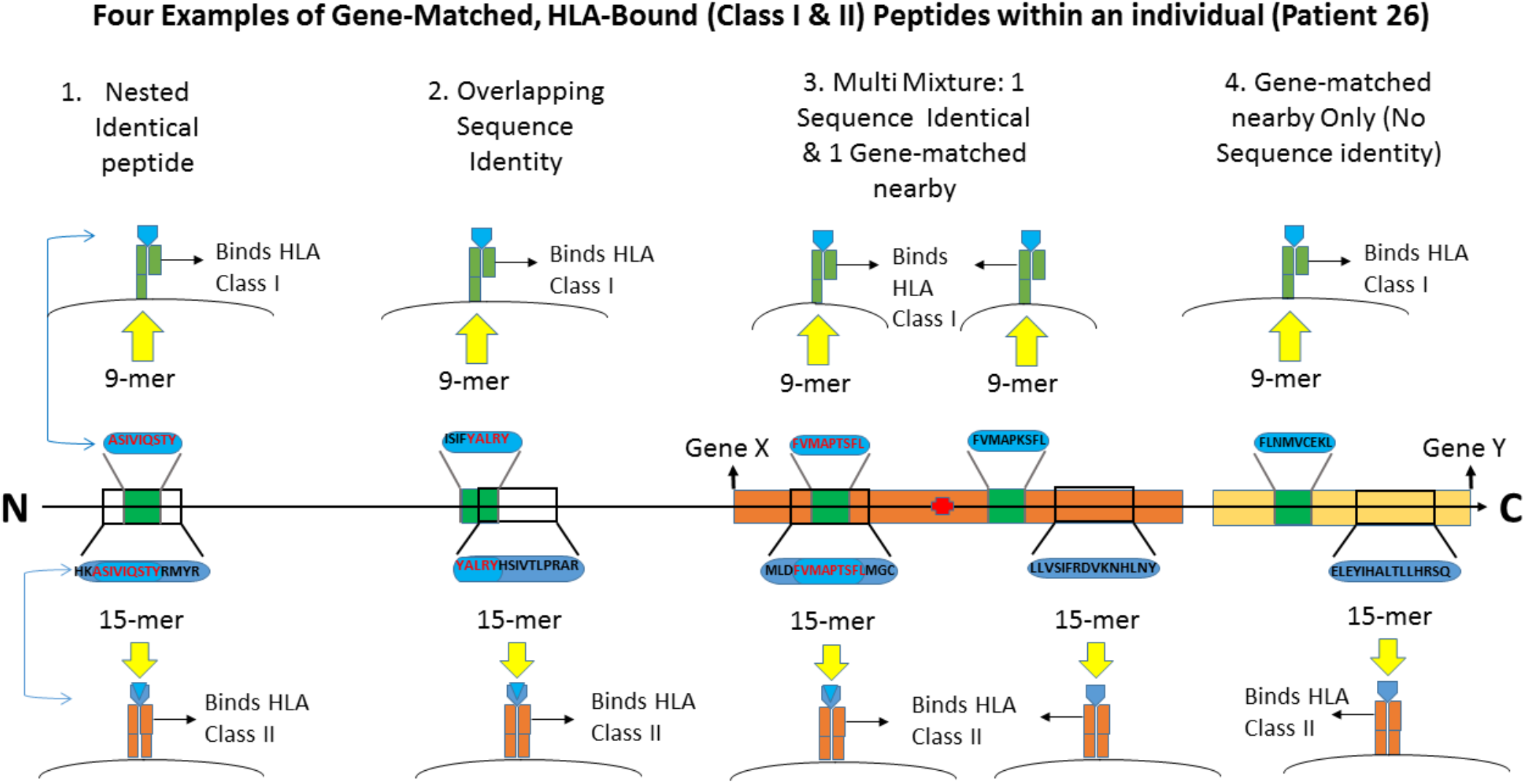
Patient 26 gene-matched, HLA-bound (class I & II) peptides with binding affinity values in the IC50, 0-50 nM range with varying degrees of sequence identity and mixtures of match types within the same gene. 1) Nested Identical peptide – completely identical amino acid sequences in the 9-mer which binds HLA Class I and the 15-mer which binds HLA Class II (peptide n=202); 2) Overlapping Sequence – overlapping amino acid sequences (3 or greater consecutively) in both 9-mer and 15-mer peptides which bind their respective HLA classes (n=268); 3) Multiple mixture: At least 1 sequence identical or nested match and at least 1 gene-matched set of discrete peptides nearby (exhibiting no sequence identity, only matched at the gene-expression level, Gene X above) (n=102); 4) gene-matched set of discrete peptides nearby to one another in proximity (exhibiting no sequence identity, only matched at the gene-expression level, Gene Y above) (n=143).

**Supplemental Figure 2.**
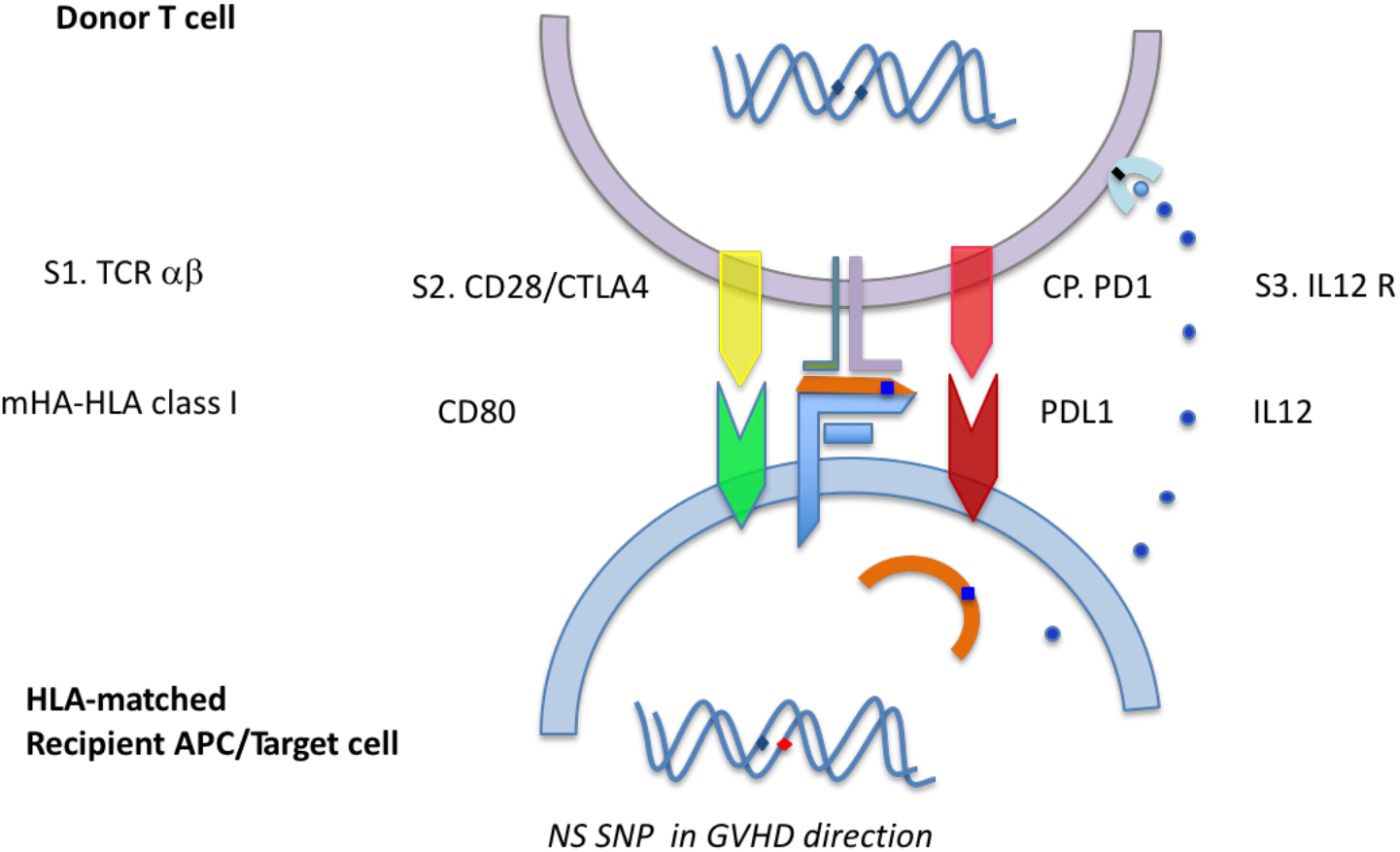
Control of T cell growth rate by the immunological synapse and Signals S1 (TCR-αβ), S2 (CD28/CTLA-4) and S3 (IL-12R), along with checkpoint molecules (CP) PD1.

**Supplemental Figure 3.**
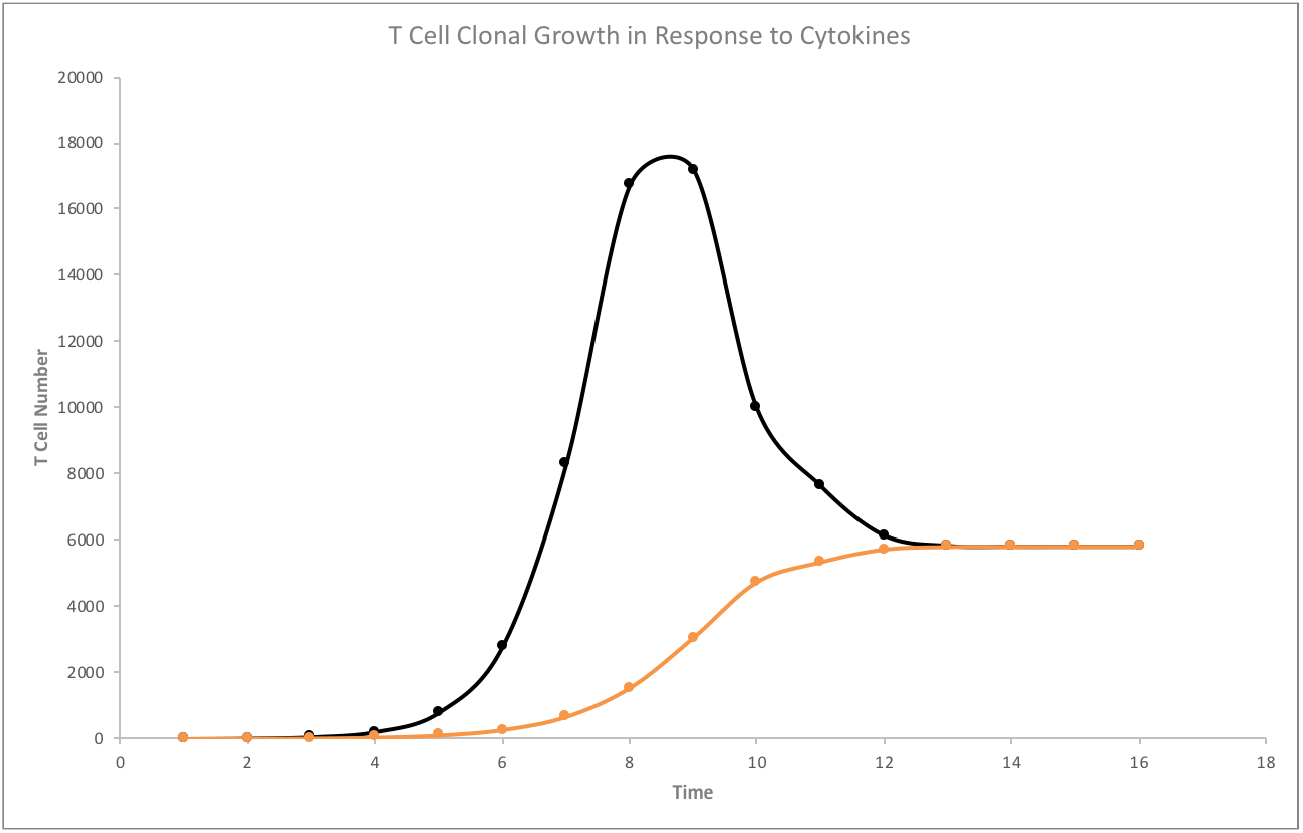
Effect of the cytokine tensor on the growth of T cells over time. Black line indicates the growth of a T cell clone with cytokine effect accounted for. Orange line depicts growth with no cytokine mediated amplification. In calculating these T cell growth trajectories, *N_0_* is 1, *K* is 1000000, *B* is 0.62 nM^-1^, *Z* is set at 1 nM^-1^, *P.c* is set at 1 RPKM, *Ck* is 1.5, *CP* & *SD* are both set at 1 (Equation 12).

## References

1. Storb R, Gyurkocza B, Storer BE, Sorror ML, Blume K, Niederwieser D, Chauncey TR, Pulsipher MA, Petersen FB, Sahebi F, Agura ED, Hari P, Bruno B, McSweeney PA, Maris MB, Maziarz RT, Langston AA, Bethge W, Vindeløv L, Franke GN, Laport GG, Yeager AM, Hübel K, Deeg HJ, Georges GE, Flowers ME, Martin PJ, Mielcarek M, Woolfrey AE, Maloney DG, Sandmaier BM. Graft-versus-host disease and graft-versus-tumor effects after allogeneic hematopoietic cell transplantation. J Clin Oncol. 2013 Apr 20;31(12):1530–8.

2. Spierings E, Kim YH, Hendriks M, Borst E, Sergeant R, Canossi A, Oudshoorn M, Loiseau P, Dolstra H, Markiewicz M, Leffell MS, Pereira N, Kircher B, Turpeinen H, Eliaou JF, Gervais T, Laurin D, Enczmann J, Martinetti M, Thomson J, Oguz F, Santarone S, Partanen J, Siekiera U, Alessandrino EP, Kalayoglu S, Brand R, Goulmy E. Multicenter analyses demonstrate significant clinical effects of minor histocompatibility antigens on GvHD and GvL after HLA-matched related and unrelated hematopoietic stem cell transplantation. Biol Blood Marrow Transplant. 2013 Aug;19(8):1244–53. doi: 10.1016/j.bbmt.2013.06.001.

3. Kosugi-Kanaya M, Ueha S, Abe J, Shichino S, Shand FHW, Morikawa T, Kurachi M, Shono Y, Sudo N, Yamashita A, Suenaga F, Yokoyama A, Yong W, Imamura M, Teshima T, Matsushima K. Long-Lasting Graft-Derived Donor T Cells Contribute to the Pathogenesis of Chronic Graft-versus-Host Disease in Mice. Front Immunol. 2017 Dec 18;8:1842. doi: 10.3389/fimmu.2017.01842.

4. Amir AL, van der Steen DM, Hagedoorn RS, Kester MG, van Bergen CA, Drijfhout JW, de Ru AH, Falkenburg JH, van Veelen PA, Heemskerk MH. Allo-HLA-reactive T cells inducing graft-versus-host disease are single peptide specific. Blood. 2011 Dec 22;118(26):6733–42.

5. Ferrara GB, Bacigalupo A, Lamparelli T, et al. Bone marrow transplantation from unrelated donors: the impact of mismatches with substitutions at position 116 of the human leukocyte antigen class I heavy chain. Blood 2001;98:3150

6. Woolfrey A, Klein JP, Haagenson M, Spellman S, Petersdorf E, Oudshoorn M, Gajewski J, Hale GA, Horan J, Battiwalla M, Marino SR, Setterholm M, Ringden O, Hurley C, Flomenberg N, Anasetti C, Fernandez-Vina M, Lee SJ. HLA-C antigen mismatch is associated with worse outcome in unrelated donor peripheral blood stem cell transplantation. Biol Blood Marrow Transplant. 2011 Jun;17(6):885–92.

7. Griffioen M, van Bergen CA, Falkenburg JH. Autosomal Minor Histocompatibility Antigens: How Genetic Variants Create Diversity in Immune Targets. Front Immunol. 2016 Mar 15;7:100.

8. Griffioen M, van Bergen CA, Falkenburg JH. Autosomal Minor Histocompatibility Antigens: How Genetic Variants Create Diversity in Immune Targets. Front Immunol. 2016 Mar 15;7:100.

9. Terakura S, Murata M, Warren EH, Sette A, Sidney J, Naoe T, Riddell SR. A single minor histocompatibility antigen encoded by UGT2B17 and presented by human leukocyte antigen-A*2902 and -B*4403. Transplantation. 2007 May 15;83(9):1242–8.

10. Vogt MH, van den Muijsenberg JW, Goulmy E, Spierings E, Kluck P, Kester MG, van Soest RA, Drijfhout JW, Willemze R, Falkenburg JH. The DBY gene codes for an HLA-DQ5-restricted human male-specific minor histocompatibility antigen involved in graft-versus-host disease. Blood. 2002 Apr 15;99(8):3027–32.

11. Abelin JG, Keskin DB, Sarkizova S, Hartigan CR, Zhang W, Sidney J, Stevens J, Lane W, Zhang GL, Eisenhaure TM, Clauser KR, Hacohen N, Rooney MS, Carr SA, Wu CJ. Mass Spectrometry Profiling of HLA-Associated Peptidomes in Mono-allelic Cells Enables More Accurate Epitope Prediction. Immunity. 2017 Feb 21;46(2):315–326.

12. Ofran Y, Kim HT, Brusic V, Blake L, Mandrell M, Wu CJ, Sarantopoulos S, Bellucci R, Keskin DB, Soiffer RJ, Antin JH, Ritz J. Diverse patterns of T-cell response against multiple newly identified human Y chromosome-encoded minor histocompatibility epitopes. Clin Cancer Res. 2010 Mar 1;16(5):1642–51.

13. Armistead PM, Liang S, Li H, Lu S, Van Bergen CA, Alatrash G, St John L, Hunsucker SA, Sarantopoulos S, Falkenburg JH, Molldrem JJ. Common minor histocompatibility antigen discovery based upon patient clinical outcomes and genomic data. PLoS One. 2011;6(8):e23217.

14. Martin PJ, Levine DM, Storer BE, Warren EH, Zheng X, Nelson SC, Smith AG, Mortensen BK, Hansen JA. Genome-wide minor histocompatibility matching as related to the risk of graft-versus-host disease. Blood. 2017 Feb 9;129(6):791–798.

15. Sampson JK, Sheth NU, Koparde VN, Scalora AF, Serrano MG, Lee V, Roberts CH, Jameson-Lee M, Ferreira-Gonzalez A, Manjili MH, Buck GA, Neale MC, Toor AA. Whole exome sequencing to estimate alloreactivity potential between donors and recipients in stem cell transplantation. Br J Haematol. 2014 Aug;166(4):566–70.

16. Pineda S, Sigdel TK, Chen J, Jackson AM, Sirota M, Sarwal MM. Novel Non-Histocompatibility Antigen Mismatched Variants Improve the Ability to Predict Antibody-Mediated Rejection Risk in Kidney Transplant. Front Immunol. 2017 Dec 5;8:1687.

17. Hoof I, Peters B, Sidney J, Pedersen LE, Sette A, Lund O, Buus S, Nielsen M. NetMHCpan, a method for MHC class I binding prediction beyond humans. Immunogenetics. 2009 Jan;61(1):1–13.

18. Rasmussen M, Fenoy E, Harndahl M, Kristensen AB, Nielsen IK, Nielsen M, Buus S. Pan-Specific Prediction of Peptide-MHC Class I Complex Stability, a Correlate of T Cell Immunogenicity. J Immunol. 2016 Aug 15;197(4):1517–24.

19. Zhang H, Lundegaard C, Nielsen M. Pan-specific MHC class I predictors: a benchmark of HLA class I pan-specific prediction methods. Bioinformatics. 2009 Jan 1;25(1):83–9.

20. Jameson-Lee M, Koparde V, Griffith P, Scalora AF, Sampson JK, Khalid H, Sheth NU, Batalo M, Serrano MG, Roberts CH, Hess ML, Buck GA, Neale MC, Manjili MH, Toor AA. In silico Derivation of HLA-Specific Alloreactivity Potential from Whole Exome Sequencing of Stem-Cell Transplant Donors and Recipients: Understanding the Quantitative Immunobiology of Allogeneic Transplantation. Front Immunol. 2014 Nov 6;5:529.

21. Granados DP, Sriranganadane D, Daouda T, Zieger A, Laumont CM, Caron-Lizotte O, Boucher G, Hardy MP, Gendron P, Côté C, Lemieux S, Thibault P, Perreault C. Impact of genomic polymorphisms on the repertoire of human MHC class I-associated peptides. Nat Commun. 2014 Apr 9;5:3600.

22. Abdul Razzaq B, Scalora A, Koparde VN, Meier J, Mahmood M, Salman S, Jameson-Lee M, Serrano MG, Sheth N, Voelkner M, Kobulnicky DJ, Roberts CH, Ferreira-Gonzalez A, Manjili MH, Buck GA, Neale MC, Toor AA. Dynamical System Modeling to Simulate Donor T Cell Response to Whole Exome Sequencing-Derived Recipient Peptides Demonstrates Different Alloreactivity Potential in HLA-Matched and -Mismatched Donor-Recipient Pairs. Biol Blood Marrow Transplant. 2016 May;22(5):850–61.

23. Koparde V, Abdul Razzaq B, Suntum T, Sabo R, Scalora A, Serrano M, Jameson-Lee M, Hall C, Kobulnicky D, Sheth N, Feltz J, Contaifer D Jr, Wijesinghe D, Reed J, Roberts C, Qayyum R, Buck G, Neale M, Toor A. Dynamical system modeling to simulate donor T cell response to whole exome sequencing-derived recipient peptides: Understanding randomness in alloreactivity incidence following stem cell transplantation. PLoS One. 2017 Dec 1;12(12):e0187771.

24. Valcárcel D, Sierra J, Wang T, Kan F, Gupta V, Hale GA, Marks DI, McCarthy PL, Oudshoorn M, Petersdorf EW, Ringdén O, Setterholm M, Spellman SR, Waller EK, Gajewski JL, Marino SR, Senitzer D, Lee SJ. One-antigen mismatched related versus HLA-matched unrelated donor hematopoietic stem cell transplantation in adults with acute leukemia: Center for International Blood and Marrow TransplantResearch results in the era of molecular HLA typing. Biol Blood Marrow Transplant. 2011 May;17(5):640–8.

25. Konuma T, Tsukada N, Kanda J, Uchida N, Ohno Y, Miyakoshi S, Kanamori H, Hidaka M, Sakura T, Onizuka M, Kobayashi N, Sawa M, Eto T, Matsuhashi Y, Kato K, Ichinohe T, Atsuta Y, Miyamura K; Donor/Source Working Group of the Japan Society for Hematopoietic Cell Transplantation. Comparison of transplant outcomes from matched sibling bone marrow or peripheral blood stem cell and unrelated cord blood in patients 50 years or older. Am J Hematol. 2016 May;91(5):E284–92.

26. Toor AA, Kobulnicky JD, Salman S, Roberts CH, Jameson-Lee M, Meier J, Scalora A, Sheth N, Koparde V, Serrano M, Buck GA, Clark WB, McCarty JM, Chung HM, Manjili MH, Sabo RT, Neale MC. Stem cell transplantation as a dynamical system: are clinical outcomes deterministic? Front Immunol. 2014 Dec 3;5:613.

27. van Niel G, Wubbolts R, Stoorvogel W. Endosomal sorting of MHC class II determines antigen presentation by dendritic cells. Curr Opin Cell Biol. 2008 Aug;20(4):437–44.

28. Diez-Rivero CM, Lafuente EM, Reche PA. Computational analysis and modeling of cleavage by the immunoproteasome and the constitutive proteasome. BMC Bioinformatics. 2010 Sep 23;11:479

29. Lai HY, Chou TY, Tzeng CH, Lee OK. Cytokine profiles in various graft-versus-host disease target organs following hematopoietic stem cell transplantation. Cell Transplant. 2012;21(9):2033–45.

30. Furlan SN, Watkins B, Tkachev V, Cooley S, Panoskaltsis-Mortari A, Betz K, Brown M, Hunt DJ, Schell JB, Zeleski K, Yu A, Giver CR, Waller EK, Miller JS, Blazar BR, Kean LS. Systems analysis uncovers inflammatory Th/Tc17-driven modules during acute GVHD in monkey and human T cells. Blood. 2016 Nov 24;128(21):2568–2579.

31. Valbon SF, Condotta SA, Richer MJ. Regulation of effector and memory CD8(+) T cell function by inflammatory cytokines. Cytokine. 2016 Jun;82:16–23.

32 Kanakry CG, Coffey DG, Towlerton AM, Vulic A, Storer BE, Chou J, Yeung CC, Gocke CD, Robins HS, O’Donnell PV, Luznik L, Warren EH. Origin and evolution of the T cell repertoire after posttransplantation cyclophosphamide. JCI Insight. 2016;1(5). pii: e86252.

33. van Heijst JW, Ceberio I, Lipuma LB, Samilo DW, Wasilewski GD, Gonzales AM, Nieves JL, van den Brink MR, Perales MA, Pamer EG. Quantitative assessment of T cell repertoire recovery after hematopoietic stem cell transplantation. Nat Med. 2013 Mar;19(3):372–7.

34. Choo SY. The HLA System: Genetics, Immunology, Clinical Testing, and Clinical Implications. Yonsei Medical Journal. 2007;48(1):11–23. doi:10.3349/ymj.2007.48.1.11

35. Bassani-Sternberg M, Pletscher-Frankild S, Jensen LJ, Mann M. Mass spectrometry of human leukocyte antigen class I peptidomes reveals strong effects of protein abundance and turnover on antigen presentation. Mol Cell Proteomics. 2015 Mar;14(3):658–73.

36. Neefjes J, Jongsma ML, Paul P, Bakke O. Towards a systems understanding of MHC class I and MHC class II antigen presentation. Nat Rev Immunol. 2011 Nov 11;11(12):823–36.

37. Meier J, Roberts C, Avent K, Hazlett A, Berrie J, Payne K, Hamm D, Desmarais C, Sanders C, Hogan KT, Archer KJ, Manjili MH, Toor AA. Fractal organization of the human T cell repertoire in health and after stem cell transplantation. Biol Blood Marrow Transplant. 2013 Mar;19(3):366–77.

38. Bolkhovskaya OV, Zorin DY, Ivanchenko MV. Assessing T cell clonal size distribution: a non-parametric approach. PLoS One. 2014 Oct 2;9(9):e108658.

39. Martínez C, Urbano-Ispizua A, Rozman C, Marín P, Rovira M, Sierra J, Montfort N, Carreras E, Montserrat E. Immune reconstitution following allogeneic peripheral blood progenitor cell transplantation: comparison of recipients of positive CD34+ selected grafts with recipients of unmanipulated grafts. Exp Hematol. 1999 Mar;27(3):561–8.

40. Pidala J, Anasetti C, Kharfan-Dabaja MA, Cutler C, Sheldon A, Djulbegovic B. Decision analysis of peripheral blood versus bone marrow hematopoietic stem cells for allogeneic hematopoietic cell transplantation. Biol Blood Marrow Transplant. 2009 Nov;15(11):1415–21.

41. Koreth J, Pidala J, Perez WS, Deeg HJ, Garcia-Manero G, Malcovati L, Cazzola M, Park S, Itzykson R, Ades L, Fenaux P, Jadersten M, Hellstrom-Lindberg E, Gale RP, Beach CL, Lee SJ, Horowitz MM, Greenberg PL, Tallman MS, DiPersio JF, Bunjes D, Weisdorf DJ, Cutler C. Role of reduced-intensity conditioning allogeneic hematopoietic stemcell transplantation in older patients with de novo myelodysplastic syndromes: an international collaborative decision analysis. J Clin Oncol. 2013 Jul 20;31(21):2662–70.

42. Hülsdünker J, Zeiser R. Insights into the pathogenesis of GvHD: what mice can teach us about man. Tissue Antigens. 2015 Jan;85(1):2–9.

43. Schroeder MA, DiPersio JF. Mouse models of graft-versus-host disease: advances and limitations. Dis Model Mech. 2011 May;4(3):318–33.

44. James Nearing. Tensors. Mathematical Tools for Physics. Pp. 327–359. Dover Publications Inc.

45. Feynman, Leighton & Sands. Tensors. The Feynman Lectures On Physics. Chapter 31 (online version). California Institute of Technology.

46. Dan Fleisch. Higher Rank Tensors. A Students Guide to Vectors and Tensors. Pp 132. Cambridge University Press.

47. Thomas Jordan. Matrices. Quantum Mechanics in Simple Matrix Form. Pp 19. Dover Publications

48. Hall CE, Koparde VN, Jameson-Lee M, Elnasseh AG, Scalora AF, Kobulnicky DJ, Serrano MG, Roberts CH, Buck GA, Neale MC, Nixon DE, Toor AA. Sequence homology between HLA-bound cytomegalovirus and human peptides: A potential trigger for alloreactivity. PLoS One. 2017 Aug 11;12(8):e0178763.

49. Pietropaolo M, Surhigh JM, Nelson PW, Eisenbarth GS. Primer: Immunity and Autoimmunity. Diabetes. 2008;57(11):2872–2882.

50. Miller DM, Thornley TB, Greiner DL, Rossini AA. Viral Infection: A Potent Barrier to Transplantation Tolerance. Clinical and Developmental Immunology. 2008;2008:742810.

51. O’Donnell JS, Smyth MJ, Teng MWL. PD1 functions by inhibiting CD28-mediated co-stimulation. Clinical & Translational Immunology. 2017;6(5):e138.

52. Bonacci B, Edwards B, Jia S, Williams CB, Hessner MJ, Gauld SB, Verbsky JW. Requirements for growth and IL-10 expression of highly purified human T regulatory cells. J Clin Immunol. 2012 Oct;32(5):1118–28.

53. Pidala J, Lee SJ, Ahn KW, Spellman S, Wang HL, Aljurf M, Askar M, Dehn J, Fernandez Viña M, Gratwohl A, Gupta V, Hanna R, Horowitz MM, Hurley CK, Inamoto Y, Kassim AA, Nishihori T, Mueller C, Oudshoorn M, Petersdorf EW, Prasad V, Robinson J, Saber W, Schultz KR, Shaw B, Storek J, Wood WA, Woolfrey AE, Anasetti C. Nonpermissive HLA-DPB1 mismatch increases mortality after myeloablative unrelated allogeneic hematopoietic cell transplantation. Blood. 2014 Oct 16;124(16):2596–606.

54. Gallardo D, Brunet S, Torres A, Alonso-Nieto M, Vallejo C, Jiménez A, González M, Sanz G, Serrano D, Espigado I, Osorio S, Carreras E, Martiín C, Sanz-Rodríguez C, Sierra J, Zuazu J, González-Escribano MF, González JR, Román J, De Oteyza JP, De La Cámara R. Hla-DPB1 mismatch in HLA-A-B-DRB1 identical sibling donor stem cell transplantation and acute graft-versus-host disease. Transplantation. 2004 Apr 15;77(7):1107–10.

55. Kekre N, Mak KS, Stopsack KH, Binder M, Ishii K, Brånvall E, Cutler CS. Impact of HLA-Mismatch in Unrelated Donor Hematopoietic Stem Cell Transplantation: A Meta-Analysis. Am J Hematol. 2016 Jun;91(6):551–5.

56. Petersdorf EW, Malkki M, O’hUigin C, Carrington M, Gooley T, Haagenson MD, Horowitz MM, Spellman SR, Wang T, Stevenson P. High HLA-DP Expression and Graft-versus-Host Disease. N Engl J Med. 2015 Aug 13;373(7):599–609.

57. Altuvia Y, Margalit H. Sequence signals for generation of antigenic peptides by the proteasome: implications for proteasomal cleavage mechanism. J Mol Biol. 2000 Jan 28;295(4):879–90.

58. Muranski P, Boni A, Wrzesinski C, Citrin DE, Rosenberg SA, Childs R, Restifo NP. Increased intensity lymphodepletion and adoptive immunotherapy–how far can we go? Nat Clin Pract Oncol. 2006 Dec;3(12):668–81.

59. Abboud R, Keller J, Slade M, DiPersio JF, Westervelt P, Rettig MP, Meier S, Fehniger TA, Abboud CN, Uy GL, Vij R, Trinkaus KM, Schroeder MA, Romee R. Severe Cytokine-Release Syndrome after T Cell-Replete Peripheral Blood Haploidentical Donor Transplantation Is Associated with Poor Survival and Anti-IL-6 Therapy Is Safe and Well Tolerated. Biol Blood Marrow Transplant. 2016 Oct;22(10):1851–1860.

60. Kriecherbauer T, Marklof J, Soshnikov A. Random matrices and quantum chaos. Proceedings of the National Academy of Sciences of the United States of America. 2001;98(19):10531–10532.

